# Convergence of Angiotensin Signaling on Lung Pericyte and Stromal Behaviors

**DOI:** 10.1101/2024.06.17.599425

**Authors:** Kynon J M Benjamin, Elizabeth Gonye, Maor Sauler, Sarah Gidner, Alla Malinina, Enid R Neptune

## Abstract

The renin-angiotensin system is a well-characterized regulator of tissue homeostasis whose clinical relevance has expanded to include lung disorders such as chronic obstructive pulmonary disease (COPD)-associated emphysema, idiopathic pulmonary fibrosis, and COVID-19. Despite this interest, the cell-specific localization of angiotensin receptors in the human lung has remained poorly defined, in part due to limitations of available antibody reagents. Here, we define the expression patterns of the two predominant angiotensin receptors, *AGTR1* and *AGTR2*, using complementary bulk and single-nucleus transcriptomic datasets from human lung tissue. We demonstrate that these receptors exhibit mutually exclusive, compartment-specific localization, with *AGTR1* expressed in lung pericytes and *AGTR2* expressed in alveolar epithelial type 2 cells. *AGTR1* is detectable in isolated lung pericytes, and spatial colocalization with pericyte markers confirmed within the airspace microvasculature compartment by RNAscope. Airspace pericyte abundance was reduced in an experimental emphysema model but restored by pharmacologic attenuation of *AGTR1* signaling commensurate with airspace repair. In COPD lungs, *AGTR1* expression showed heterogeneous, disease-associated dysregulation across stromal populations, including upregulation in alveolar fibroblasts. Bulk transcriptomics also revealed aging-associated redistribution of *AGTR1* expression into stromal compartments. Angiotensin II and cigarette smoke impaired pericyte migration toward endothelial cells, while combined exposure suppressed pericyte proliferation. Together, these findings identify *AGTR1* as a new highly selective marker of lung pericytes and a regulator of pericyte behaviors within the airspace microvasculature. These findings provide a cell-resolved framework for angiotensin signaling with direct relevance to airspace resilience and therapeutic targeting.

## Introduction

The renin-angiotensin system (RAS) is an extensively characterized hormonal network that plays a central role in maintaining mammalian tissue homeostasis. Dysregulated RAS signaling contributes to numerous common disorders, including hypertension, chronic renal disease, and heart failure^1^, supporting the widespread clinical use of RAS-targeting therapies. Indeed, angiotensin receptor blockers (ARBs) and angiotensin-converting enzyme inhibitors are consistently among the most prescribed medications in the United States^2–8^.

Increasing evidence has reframed RAS activity as highly tissue- and context-dependent, with roles extending beyond systemic hemodynamic regulation. In the lung, angiotensin signaling has been implicated in inflammation and tissue repair^9^, acute lung injury^10,11^, idiopathic pulmonary fibrosis (IPF)^12^, pulmonary hypertension^13,14^, and chronic obstructive pulmonary disease (COPD)-associated emphysema^15,16^. The identification of ACE2 as the SARS-CoV-2 receptor further highlighted the relevance of this pathway to lung disease pathogenesis^17,18^.

Despite this broad interest, a clear delineation of the abundance and cell-specific expression pattern of angiotensin receptors in the human lung remains lacking.

One major barrier to defining angiotensin receptor localization has been the limited specificity of available antibody reagents, a limitation that has been insufficiently appreciated in prior studies^19–21^. Consequently, inaccurate localization data have contributed to conceptual ambiguity regarding lung cell-specific receptor function and, potentially, to variable outcomes in preclinical and clinical trials of ARBs. Although ARB-mediated attenuation of airspace enlargement has been demonstrated in genetic^22,23^ and cigarette smoke-induced^24,25^ mouse models of emphysema, and beneficial effects have been reported in fibrotic lung disease models^26,27^, clinical efficacy has been inconsistent^8,28–30^. Beyond species differences and dosing constraints, an incomplete understanding of lung cell-specific angiotensin receptor biology likely limits rational therapeutic targeting.

In this study, we define the expression pattern of the two predominant angiotensin receptors in the human lung, *AGTR1* and *AGTR2*, using complementary bulk and single-nucleus transcriptomic datasets. We identify near-exclusive expression of *AGTR1* in lung pericytes, implicating a mechanistic role for angiotensin signaling within the microvascular compartment of the airspace. We validate *AGTR1* expression in isolated human and mouse pericytes and demonstrate that pericyte abundance and behavior are altered in experimental emphysema models in an *AGTR1*-dependent manner. By integrating human transcriptomic discovery with *in vivo* and *in vitro* functional studies, our findings establish a cell-resolved framework for angiotensin signaling in the lung and provide a mechanistic framework for understanding context-dependent responses to angiotensin-targeting therapies in chronic lung disease.

## Results

### Angiotensin receptors show distinct and mutually exclusive localization in the human lung

To identify the cellular localization of angiotensin receptors, we analyzed two human lung single-cell datasets: the Human Lung Cell Atlas (HLCA, version 2)^31^ and LungMAP^32^. The HLCA (core model: 584,944 cells from 14 datasets and 107 individuals^31^) served as the primary dataset, and we used LungMAP (46,500 cells, 9 individuals^32^) as an independent replication set after harmonizing cell annotations to the HLCA reference. Across both datasets, cells expressing angiotensin receptors 1 (*AGTR1*) or 2 (*AGTR2*) were rare, representing 1.6% of cells in HLCA and 3.0% in LungMAP. Of these receptor-positive cells, expression was highly mutually exclusive, with fewer than 0.1% co-expressed both *AGTR1* and *AGTR2* (**Figure 1A** and **Figure S1A**). We also found that *AGTR1*- and *AGTR2*-positive cells localized to distinct lung compartments (**Figure 1B** and **Figure S1B**).

**Figure 1:**
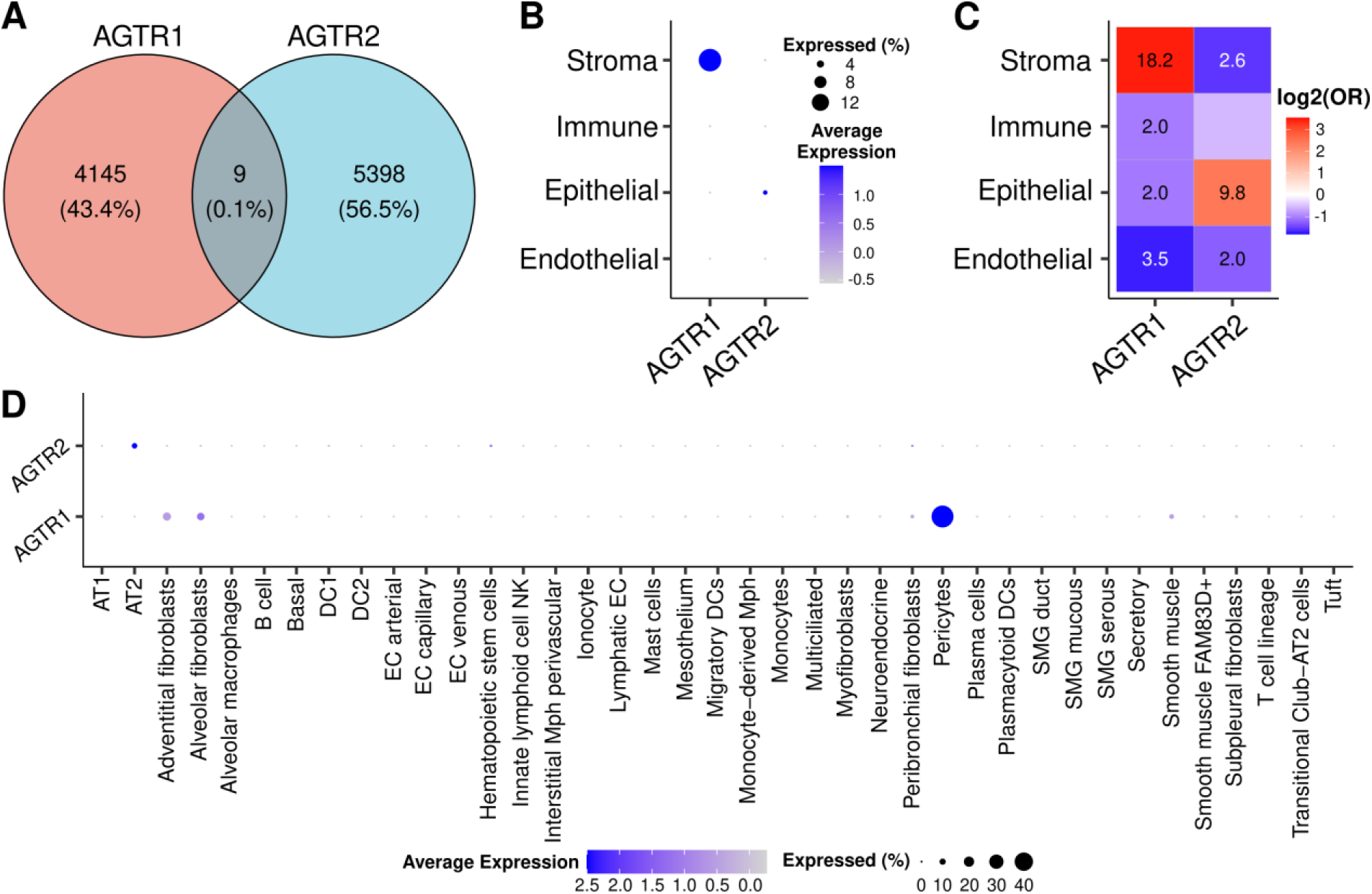
Angiotensin II receptors 1 (AGTR1) and 2 (AGTR2) demonstrate compartment-and cell-type-specific expression in the human lung. **A.** Venn diagram showing rare occurrences of *AGTR1* and *AGTR2* co-expression (HLCA^31^; n=107 individuals). B. Dot plot showing the percentage and average expression of *AGTR1*- and *AGTR2*-positive cells across lung compartments (n=107). C. Heatmap showing significant enrichment (two-sided, Fisher’s exact test) of *AGTR1*- and *AGTR2*-positive cells across lung compartments (n=107). The color intensity of enrichment heatmaps represents log2 of odds ratio (OR) with red indicating enrichment and blue indicating depletion. Significantly enrichment compartments are annotated with -log10(false discovery rate). D. Dotplot showing the percentage and average expression of *AGTR1*- and *AGTR2*-positive cells across lung cell-types (n=107).

*AGTR1*-positive cells predominated in the stromal/mesenchymal lung compartment (Fisher’s exact test, FDR < 0.05; **Figure 1C** and **Figure S1C**). Within this compartment, pericytes represented the largest fraction of *AGTR1*-expressing cells in HLCA (48% of the *AGTR1*⁺ population; Fisher’s exact test, FDR < 0.05; **Figure 1D**) and among the highest in LungMAP (16%; Fisher’s exact test, FDR < 0.05; **Figure S1D**). Other stromal cell-types, including alveolar fibroblasts (13% in HLCA, 16% in LungMAP) and adventitial fibroblasts (15% in HLCA, 11% in LungMAP), also showed significant enrichment. The larger HLCA dataset further revealed significant enrichment in vascular smooth muscle (6.4%), and peribronchial fibroblasts (4.0%) for *AGTR1*⁺ cells (Fisher’s exact test, FDR < 0.05; **Figure 1D**).

In contrast, *AGTR2*-positive cells localized primarily to the lung epithelial compartment (**Figure 1C**). The highest enrichment occurred in alveolar epithelial type 2 (AT2) cells, which comprised 8.4% of the *AGTR2*-positive population. Additionally, alveolar macrophages comprised 0.048% of *AGTR2*-positive cells (Fisher’s exact test, FDR < 0.05; **Figure 1D**). We based these *AGTR2* localization findings exclusively on the HLCA dataset because the LungMAP dataset contained an extremely low cell count of *AGTR2*-positive cells (n=51) after cell annotation harmonization. Overall, these data reveal a clear, mutually exclusive expression pattern: *AGTR1* localized to the mesenchymal stromal compartment, with pericytes as the primary expressing cell-type, while *AGTR2* localizes to the epithelial compartment, with enrichment in AT2 cells. Given the known therapeutic role of *AGTR1* antagonism in experimental emphysema, our finding that pericytes are the dominant *AGTR1*-expressing population in the lung led us to next investigate the specific role of these cells in the airspace compartment.

### AGTR1 is expressed in primary lung pericytes within the airspace compartment

Limited to nonexistent information exists concerning angiotensin signaling in pericytes – a multipotent cell-type residing in the microvascular bed and implicated in vascular development and hemodynamic regulation^33,34^. Because marker- and ontogeny-defined lung pericyte subtypes remain incompletely resolved, we first validated *AGTR1* expression in isolated primary lung pericytes. We detected high levels of *AGTR1* expression in primary pericytes isolated from human and mouse lungs, and *AGTR1* colocalized with canonical pericyte markers in the mouse airspace compartment (**Figure 2A,B**). PDGFRβ- and NG2-expressing pericytes exhibited close spatial association with CD31-expressing endothelial cells within the lung airspace (**Figure 2C**).

**Figure 2:**
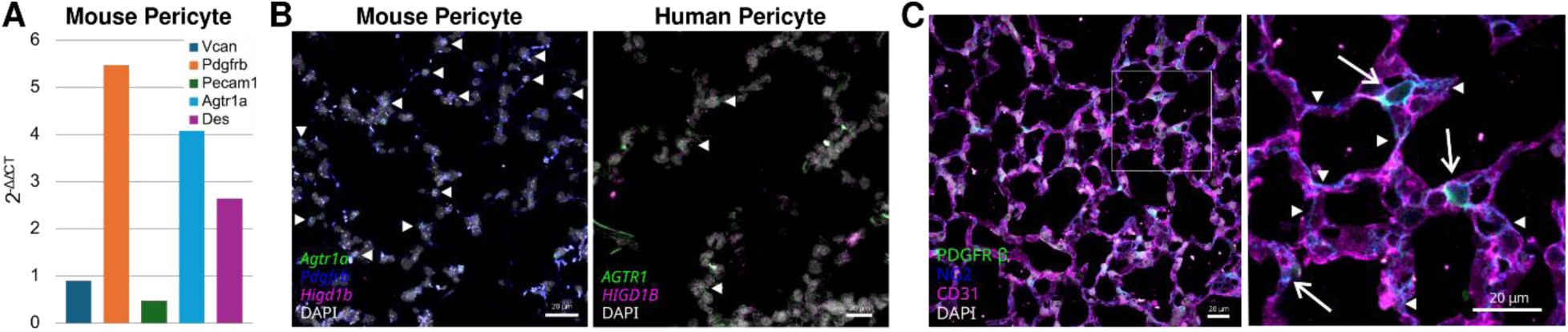
Validation of AGTR1 expression and spatial localization in lung pericytes. **A.** Isolated mouse lung pericytes express AGTR1 together with canonical pericyte markers. **B.** Representative RNAscope images showing AGTR1 localization within mouse and human lung airspace compartment and co-expression with pericyte markers NG2 and PDGFRβ. Arrows indicate cells co-expressing AGTR1 and pericyte markers. **C.** Representative immunohistochemical images showing NG2- and PDGFRβ-positive pericytes in close spatial association with CD31-expressing microvascular endothelial cells. Arrowheads denote marker-defined pericytes; inset highlights the close apposition of pericytes to microvascular endothelium. Scale bars: left, 10*μ*m; right, 20*μ*m.

### *AGTR1* expression spans transcriptionally defined lung pericyte states

Because lung pericytes comprise transcriptionally distinct subtypes and potential functional states^35^, we asked whether *AGTR1* expression in the HLCA full dataset^31^ (∼2.2 million total cells) was associated with specific pericyte classes or traditional states. Analysis of 11,680 pericytes from 194 donors identified eight transcriptionally distinct pericyte subclusters (**Figure 3A** and **Figure S2**). We detected *AGTR1* expression across subclusters (**Figure 3B**). After excluding donors with missing covariate data and subsequently restricting the analysis to clusters with at least three remaining donors, we evaluated *AGTR1* expression across four pericyte subclusters (clusters 0-3) using a linear model controlling for sex, disease status, age, and self-reported ethnicity. *AGTR1* expression differed significantly across pericyte clusters (ANOVA, *F* = 7.26, *p* = 2.4 × 10^−4^), whereas sex, disease status, and age were not associated with *AGTR1* levels; ethnicity showed a modest independent effect (ANOVA, *F* = 3.84, *p* = 0.013).

**Figure 3:**
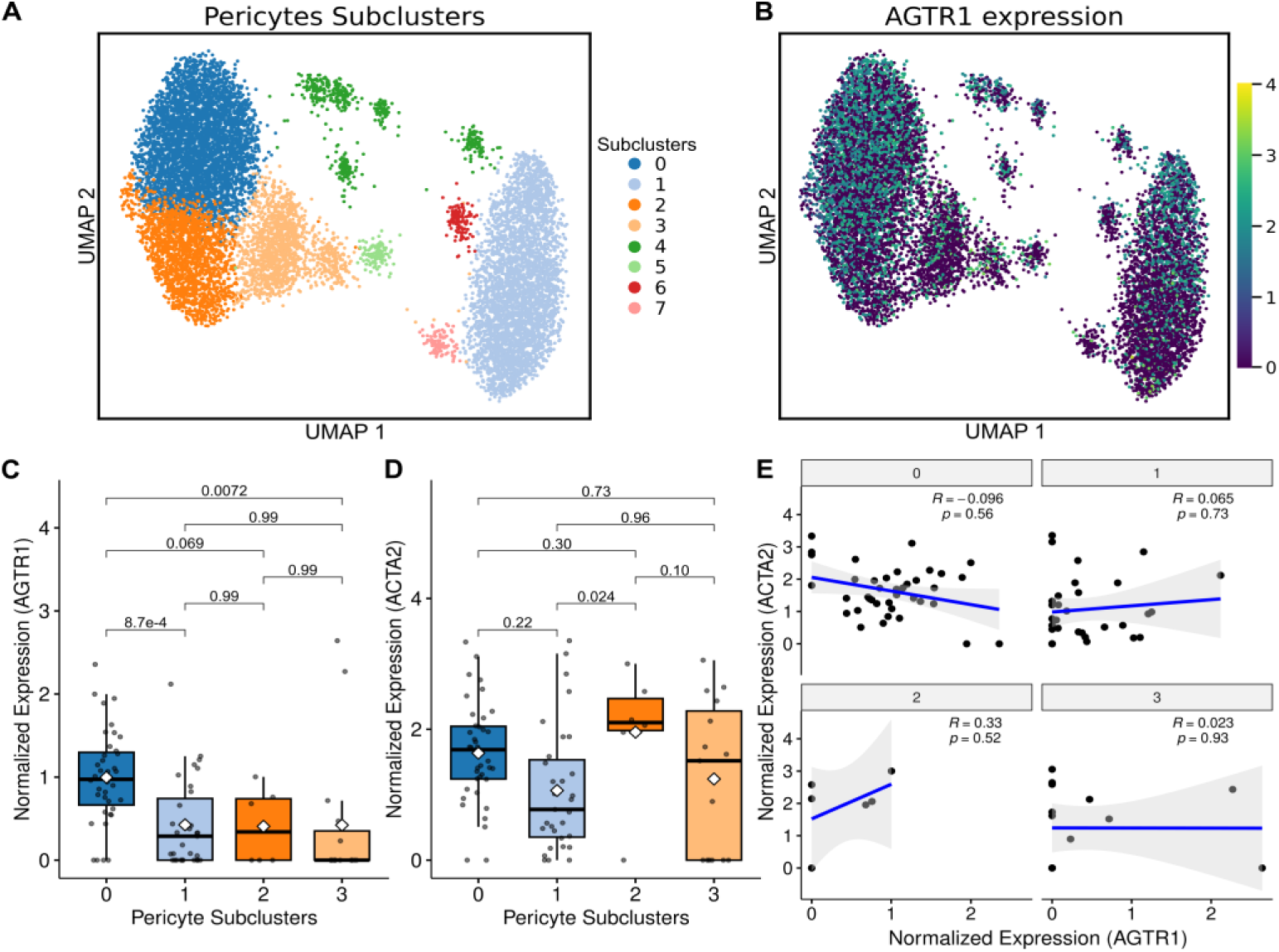
AGTR1 expression across transcriptionally defined lung pericyte subclusters. ***A.*** UMAP visualization of 11,680 lung pericytes from the HLCA full dataset^31^, colored by Leiden subcluster assignment (clusters 0-7). **B.** UMAP showing normalized AGTR1 expression across pericyte subclusters. Boxplots of normalized **C.** AGTR1 and **D.** ACTA2 expression across pericyte subclusters meeting quality-control criteria (clusters 0-3; ≥3 donors per cluster). Boxes indicate the median and interquartile range; whiskers extend to 1.5× the interquartile range; white diamonds denote the mean. Adjusted *p* values for pairwise comparisons (Tukey’s post-hoc test) are indicated. **E.** Scatter plot showing the relationship between normalized *AGTR1* and *ACTA2* expression across pericyte subclusters. Spearman’s correlation coefficient (ρ) and corresponding *p* value are indicated. The fitted trend line represents the mean ± s.d. (shaded).

Estimated marginal means indicated higher *AGTR1* expression in cluster 0 relative to clusters 1 and 3 (Tukey’s: cluster 0 vs 1, *p* = 8.7 × 10^−4^; cluster 0 vs 3, *p* = 0.007), with no significant differences among clusters 1–3, or between cluster 0 and 2 (**Figure 3C**). Thus, *AGTR1* expression shows modest quantitative enrichment in a single pericyte cluster but is detected across multiple transcriptional states, suggesting that *AGTR1* is not restricted to a discrete pericyte subtype.

In contrast, expression of the contractile marker *ACTA2* also varied across pericyte clusters (ANOVA, *F* = 3.47, *p* = 0.020), with post hoc analysis revealing significantly higher *ACTA2* expression in cluster 2 compared with cluster 1 (Tukey’s, *p* = 0.024). No other pairwise comparisons reached significance (**Figure 3D**). Notably, clusters exhibiting elevated *ACTA2* expression did not correspond to those with higher *AGTR1* expression (**Figure 3E**), indicating that *AGTR1* variation is not aligned with the canonical contractile axis of pericyte differentiation.

Although *AGTR1* was broadly expressed, a subset of pericytes lacked detectable *AGTR1* transcripts, raising the possibility of either biological heterogeneity or technical dropout. To distinguish between these alternatives, we compared *AGTR1*-positive (*AGTR1*⁺) and *AGTR1*-negative (*AGTR1*⁻) pericytes using an airspace affinity score. A linear mixed-effects model controlling for donor identity revealed no significant difference in airspace affinity between *AGTR1*⁺ and *AGTR1*⁻ pericytes (*β* = -0.002, *SE* = 0.008, *p* = 0.79; **Figure S3A,B**).

To formally evaluate whether *AGTR1*⁻ pericytes could be explained by stochastic dropout, we modeled per-cell expected zero probabilities. Among 3,032 pericytes, 1,572 lacked detectable *AGTR1* expression, a frequency indistinguishable from technical expectation (expected = 1,607; observed-to-expected ratio = 0.978; *p* = 0.89, Poisson-binomial FFT model). Consistent with technical dropout, scVI-imputed *AGTR1* expression among *AGTR1*⁻ pericytes showed no association with airspace affinity (Spearman’s *ρ* = -0.046, *p* = 0.53; **Figure S3C**). Together, these findings indicate that apparent *AGTR1*⁻ pericytes do not represent a spatially or transcriptionally distinct subpopulation but instead reflect under calling due to technical limitations of single-nucleus RNA sequencing.

### *AGTR1* expression in the context of the aging lung

Given the increased prevalence of several airspace disorders with age^36^, we next examined whether donor age was associated with *AGTR1* expression from mid- to late-life (age range = 21-81 years) using the HCLA core dataset^31^. Across top five *AGTR1*-enriched populations (pericytes, alveolar fibroblasts, adventitial fibroblasts, smooth muscle cells, and peribronchial fibroblasts), we observed no statistically significant monotonic association between age and donor-mean *AGTR1* expression after multiple-testing correction (Spearman correlation, all FDR > 0.28; **Figure 4A**). A nominal positive correlation was observed in smooth muscle cells (ρ = 0.36, *p* = 0.057), but this trend did not remain significant following FDR adjustment.

**Figure 4:**
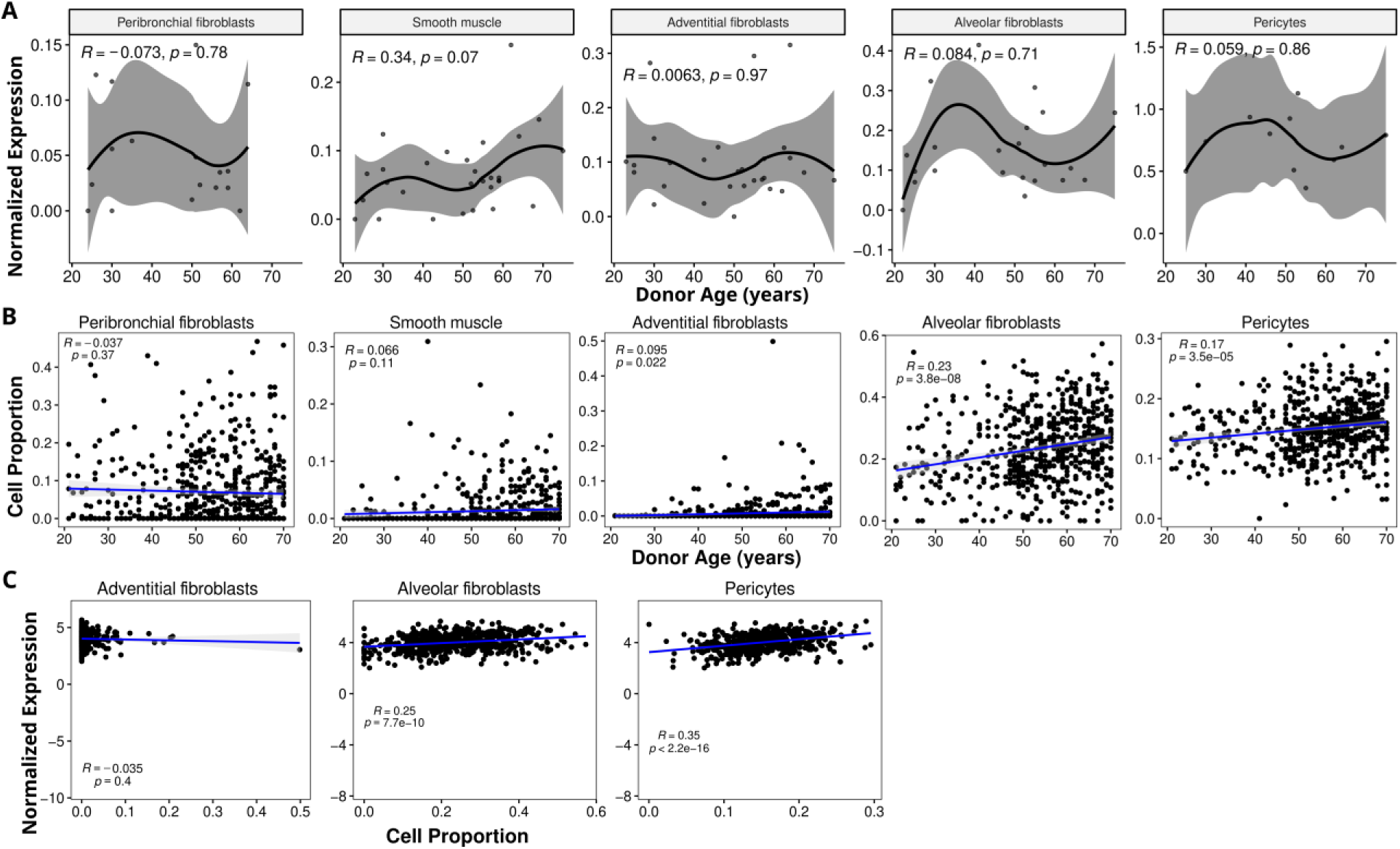
Age-associated changes in *AGTR1* expression reflect shifts in lung cellular composition. **A.** Donor-level associations between age and mean *AGTR1* expression in the HLCA core dataset^31^ (age 21-81 years) for the five most *AGTR1*-enriched populations (pericytes, alveolar fibroblasts, adventitial fibroblasts, vascular smooth muscle cells, and peribronchial fibroblasts). Each point represents one donor; the curve shows a LOESS fit.

To assess potential non-linear age effects, we next fit natural cubic spline models and generalized additive models for each cell-type. Spline-based model comparisons did not provide evidence that non-linear age terms significantly improved model fit over linear age effects for any cell-type after multiple-testing correction (all spline vs. linear FDR ≥ 0.27). Consistent with these results, GAM analyses yielded effective degrees of freedom (≈ 1) across all cell-types, indicating predominantly linear age relationships, with no significant smooth terms after FDR correction (all FDR ≥ 0.74).

Given the limited donor representation after quality control in the single-cell data, particularly pericytes (n = 11), we next extended our analysis to the GTEx v8 bulk lung RNA-sequencing dataset (age range = 21–70 years; n = 578; median age = 56)^37^. In contrast to the single-cell donor-level results, bulk lung *AGTR1* expression showed a modest but statistically significant positive association with age (Pearson’s *r* = 0.12, *p* < 0.0052).

To determine whether this bulk signal reflected age-associated changes in cellular composition, we examined inferred proportions of *AGTR1*-enriched cell types. The proportion of pericytes – the cell-type most enriched for *AGTR1*⁺ cells – was positively correlated with age (Pearson’s *r* = 0.17, *p* = 3.5 × 10^−5^; **Figure 4B**). Extending this analysis to additional *AGTR1*-enriched populations revealed positive age associations for adventitial fibroblasts and alveolar fibroblasts (Pearson’s *r* = 0.095 and 0.23; *p* = 0.022 and 3.8 × 10^−8^, respectively; **Figure 4B**). Moreover, normalized bulk *AGTR1* expression increased with the relative abundance of pericytes and alveolar fibroblasts (Pearson’s *r* = 0.35 and 0.25; *p* < 2.2 × 10^−16^ and *p* = 7.7 × 10^−10^, respectively; **Figure 4C**).

Together, these findings indicate that while *AGTR1* expression within individual lung mesenchymal cell-types is largely stable across adulthood at the donor level, age-associated increases in bulk lung *AGTR1* expression are driven primarily by shifts in the relative abundance of pericytes and alveolar fibroblasts.

Spearman correlations (*ρ*) with FDR adjustment are indicated; no cell-type remained significant after multiple-testing correction (all FDR > 0.28). **B.** Associations between age and inferred cell-type proportions in GTEx v8 bulk lung^37^ (n = 578; age 21-70 years) for *AGTR1*-enriched populations. Points represent donors; the line shows a linear regression fit. Pearson’s correlation coefficient (*r*) and *p* value are shown. **C.** Bulk lung *AGTR1* expression in GTEx versus inferred cell-type abundance, demonstrating higher *AGTR1* expression with increasing proportions of pericytes and alveolar fibroblasts. Points represent donors; the line shows a linear regression fit. Pearson’s *r* and *p* value are shown. Shaded bands indicate the 95% confidence interval.

### Disease- and cell type-specific *AGTR1* dysregulation across human lung stromal populations

To define the stromal contexts in which *AGTR1* is dysregulated in human lung disease, we examined *AGTR1* expression across stromal populations using the full HLCA dataset^31^.

Because angiotensin II signals through both *AGTR1* and *AGTR2*, pharmacologic inhibition of *AGTR1* could shift angiotensin signaling toward compensatory pathways, potentially explaining aspects of therapeutic efficacy^38^. We therefore asked whether disease-associated *AGTR1* dysregulation is restricted to specific stromal cell types.

Among the five *AGTR1*-enriched stromal populations, peribronchial fibroblasts showed the strongest and most consistent disease-associated signal. In this population, *AGTR1* expression varied significantly by disease states (Kruskal-Wallis *p* ≈ 0.0014; **Figure 5A**; **Table S1**), with multiple Dunn’s post hoc contrasts remaining significant after FDR correction. *AGTR1* expression was higher in chronic rhinitis, pulmonary fibrosis, and interstitial lung disease than in normal and COVID-19 donors, with small-to-moderate effect sizes (**Table S1**). In contrast, pericytes, vascular smooth muscle cells, alveolar fibroblasts, and adventitial fibroblasts exhibited only nominal or trend-level disease effects (overall *p* ≈ 0.07-0.09; **Table S1**), with small effect sizes and no post hoc comparisons surviving multiple-testing correction. These findings indicate that disease-associated *AGTR1* dysregulation within the stromal compartment is most pronounced in peribronchial fibroblasts.

**Figure 5:**
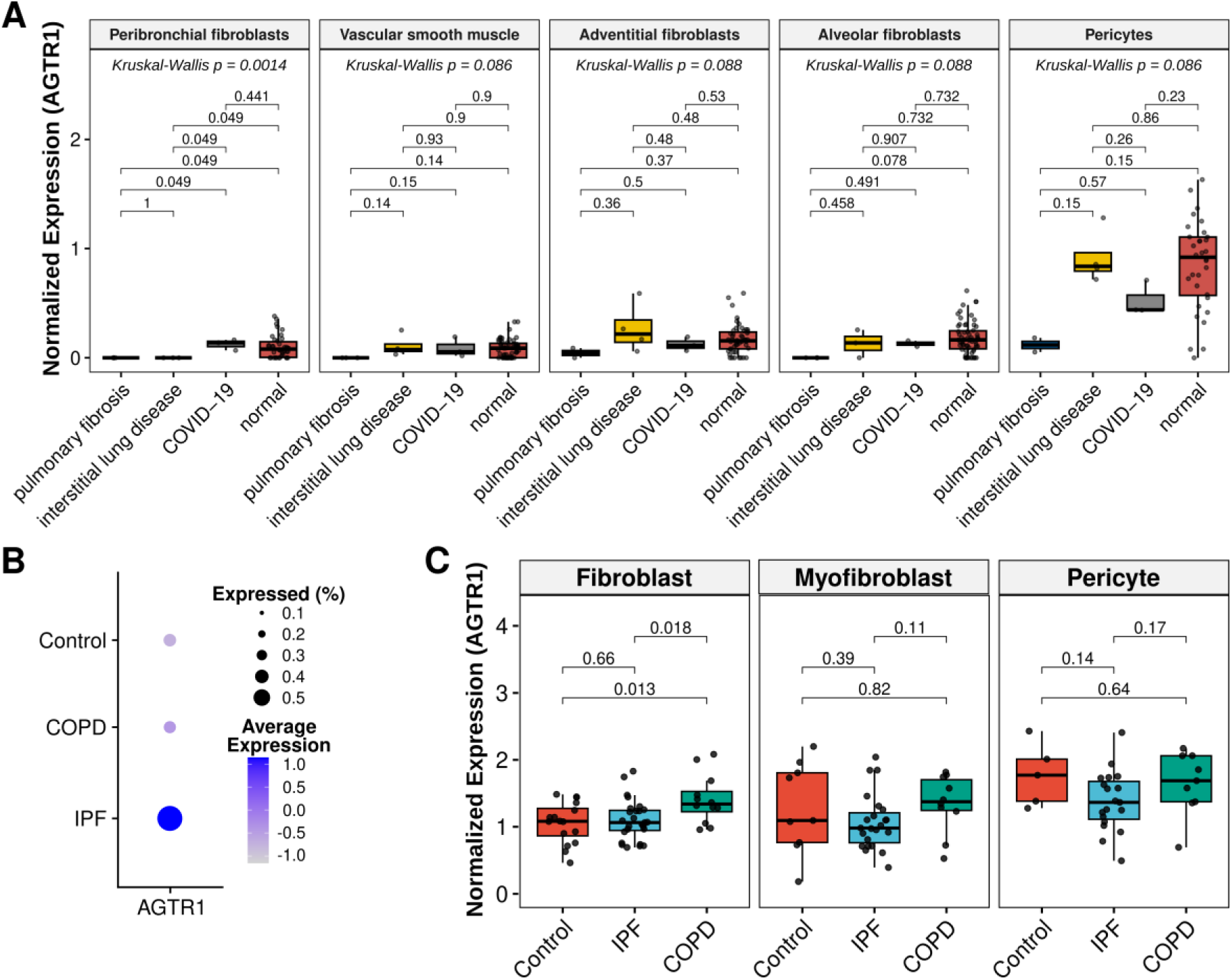
Disease-associated *AGTR1* expression across lung stromal cell types. **A.** Boxplots showing *AGTR1* expression in enriched stromal cell types across donor disease states in the HLCA dataset^31^. **B.** Dotplot of average expression and percent expression of *AGTR1* from total cells in an independent single-nucleus lung dataset^39,40^, separated by diagnosis (control [n=28], COPD [n=18], or IPF [n=32]). **C.** Boxplots from the same independent dataset showing *AGTR1* expression in *AGTR1*-enriched cell types from COPD, IPF, and control donors. All data are shown as log-normalized expression values. Boxplots display median, interquartile range (IQR), and 1.5× IQR whiskers; individual points represent donor means.

Given the modest disease signals detected in pericytes, we further examined *AGTR1* expression across transcriptionally defined pericyte subclusters to identify potential subtype-specific effects. Although no subcluster-level disease differences reached significance after multiple-testing correction, subcluster 0 exhibited a nominal overall disease effect (Kruskal-Wallis *p* ≈ 0.082; **Figure S4A**; **Table S2**). Consistent with this trend, Dunn’s post hoc tests suggested higher *AGTR1* expression in COVID-19 relative to normal donors, with moderate-to-large effect sizes despite non-significant adjusted *p*-values. Although limited by statistical power, these findings point to a potentially meaningful disease-associated signal in a particular pericyte subpopulation that warrants further investigation in larger cohorts.

Because pericytes from COPD and IPF were underrepresented in the HLCA dataset, we performed a targeted secondary analysis using an independent lung single-nucleus transcriptomic dataset enriched for these diseases, comprising IPF (n = 32), COPD (n = 18), and control donor lungs (n = 28)^39,40^. In this cohort, *AGTR1*⁺ cells were more frequently observed in IPF donors (**Figure 5B**). *AGTR1* expression was also significantly higher in fibroblasts from COPD patients compared with controls (t test, *p* < 0.013; **Figure 5C**), supporting disease-associated upregulation within mesenchymal compartments. Although mean *AGTR1* expression across all cells was highest in IPF donors, individual cell types did not differ significantly between IPF and control donors. Notably, however, a greater proportion of IPF donors (94%) harbored *AGTR1*⁺ cells compared to control (75%) or COPD (78%) donors, suggesting broader – but not cell-type-restricted – engagement of *AGTR1* in IPF.

To determine whether transcriptional heterogeneity within pericytes masked disease effects in this cohort, we subclustered pericytes using HLCA-derived annotations. No global disease effects remained significant after multiple-testing correction (**Figure S4B**; **Table S3**), although subcluster 1 showed a trend toward lower *AGTR1* expression in IPF relative to controls (Dunn’s post hoc *p* ≈ 0.039; FDR = 0.12). All other pericyte subclusters showed no evidence of significant disease-associated *AGTR1* dysregulation, with trend-level differences and small-to-moderate effect sizes (**Table S3**). Statistical power was likely constrained by small pericyte sample size (659 cells across four of eight subclusters after quality control).

Together, these analyses reveal that *AGTR1* dysregulation in human lung disease is concentrated within distinct stromal compartments – most prominently peribronchial fibroblasts – and highlight *AGTR1* signaling as a candidate pathway for functional testing in emphysema models.

### AGTR1-dependent regulation of pericyte abundance in experimental emphysema

Because relative abundance of lung cell types cannot be reliably inferred from single-cell or single-nucleus transcriptomic datasets, and *AGTR1* signaling contributes to vascular remodeling and fibrotic responses^10^, we next tested whether emphysematous injury alters pericyte abundance *in vivo* in an *AGTR1*-dependent manner. We previously reported that angiotensin receptor blockade prevented the development of chronic cigarette smoke (CS)-induced murine emphysema and reversed established emphysema in a Fbn1-variant knock-in model of Marfan Syndrome^22,24^. We explored whether the inhibition of AGTR1 signaling conferred alterations of pericyte abundance that aligned with airspace resilience in the chronic CS model. Consistent with our prior work demonstrating airspace enlargement, increased lung compliance, and oxidative injury following four months of cigarette smoke exposure in AKR/J mice, we observed an approximately 30% reduction in airspace pericyte abundance, assessed by both absolute counts and proportional representation (**Figure 6A-E**). This reduction persisted after normalization to endothelial cell investment using CD31 staining, indicating a true loss of pericytes rather than proportional remodeling. Given our previous finding that angiotensin receptor blockade prevents airspace enlargement and attenuates lung injury in chronic cigarette smoke-exposed mice, we next examined pericyte abundance following losartan treatment and found restoration of both absolute pericyte numbers and pericyte-endothelial ratios to control levels (**Figure 6E**).

**Figure 6:**
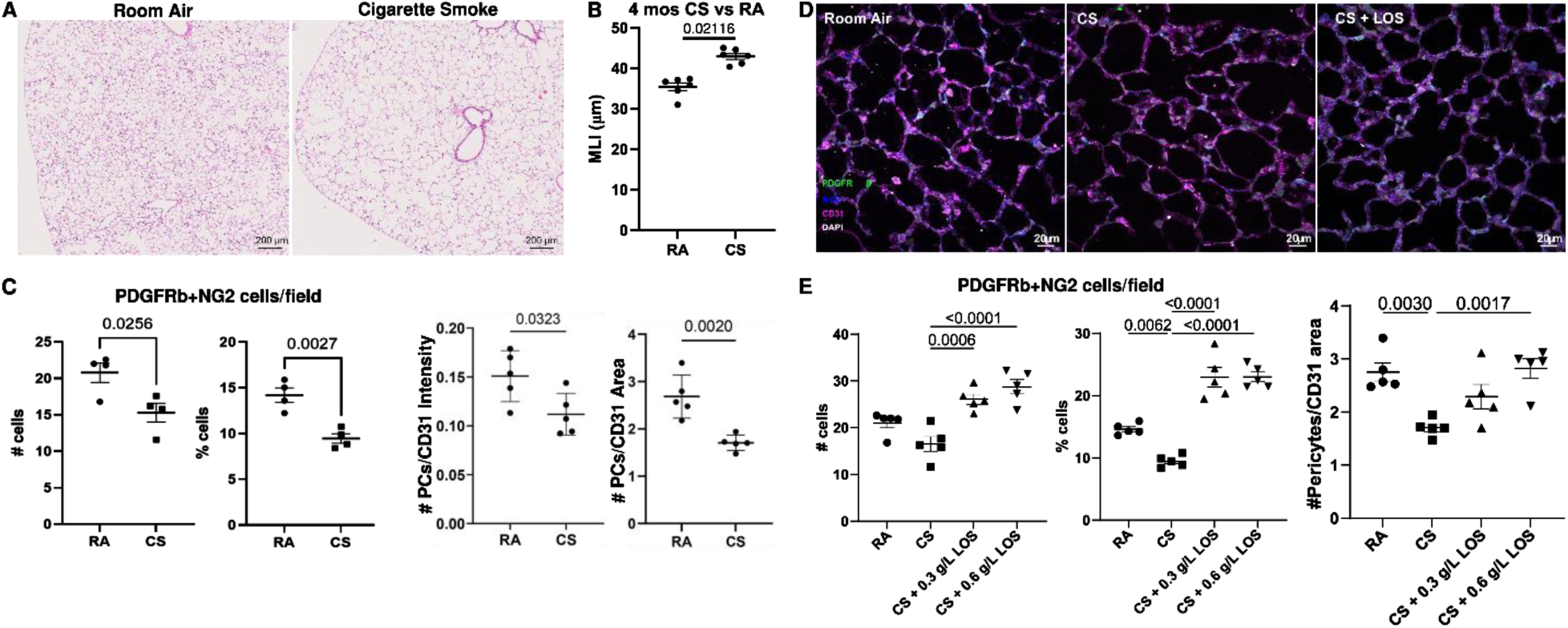
Pericyte abundance and spatial distribution in CS-induced emphysema model. **A.** Representative lung histology of mice subjected to the indicated exposures. **B.** Morphometric assessment of airspace dimensions in mice exposed to 4 months of cigarette smoke (CS) or room air (RA). **C.** Quantification of pericyte abundance in RA- and CS-exposed mice, shown as absolute cell counts, percentage of total cells, and pericyte counts normalized to endothelial cell investment assessed by CD31 staining. **D.** Representative immunofluorescent co-staining for pericytes (PDGFRβ, NG2), vascular bed (CD31), and nuclei (DAPI). Circles denote pericytes closely associated with the vascular bed, whereas arrows indicate pericytes displaced from the vascular bed in CS-exposed mice. **E.** Quantification of pericyte and endothelial cell measures in CS-exposed mice treated with two doses of losartan (LOS). *P* values are indicated above data points where applicable.

### Distinct and synergistic effects of cigarette smoke and AGTR1 activation on pericyte behaviors

Pericyte recruitment to and interaction with endothelial cells is essential for stable vascular attachment and mediation of protective functions. To determine whether *AGTR1* activation and cigarette smoke exposure independently or synergistically disrupt pericyte behaviors relevant to pericyte loss in emphysema, we treated primary human pericytes with angiotensin II (AngII) and/or cigarette smoke extract as an *in vitro* surrogate for COPD/emphysema-associated injury. Using an endothelial-pericyte transwell co-culture system, we examined candidate behaviors implicated in pericyte depletion observed in *in vivo* emphysema models.

Both cigarette smoke exposure and AngII treatment significantly inhibited pericyte migration toward endothelial cells (**Figure 7A,B**). In contrast, neither cigarette smoke nor AngII alone altered pericyte proliferation; however, combined exposure results in a marked reduction in pericyte proliferation, consistent with a synergistic inhibitory effect (**Figure 7C**).

**Figure 7:**
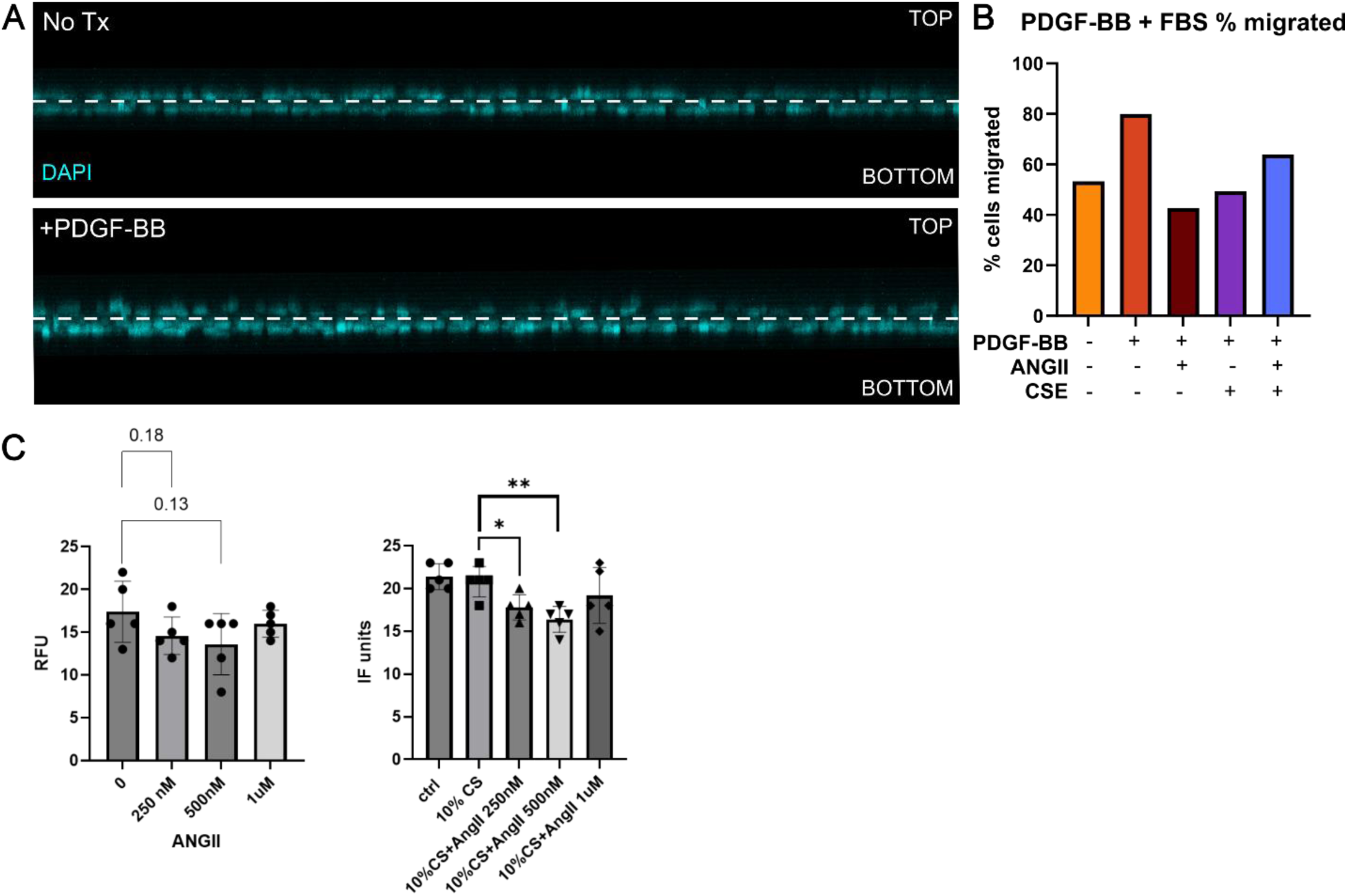
Effects of angiotensin II and cigarette smoke extract on pericyte behaviors. **A.** Representative images from a transwell migration assay showing PDGF-BB-stimulated human pericyte migration. Pericytes were seeded in serum-free medium in the upper chamber, with (+PDGF-BB) and without (No Tx) PDGF-BB and 10%FBS fetal bovine serum (FBS) in the lower chamber as a chemoattractant. The dashed line indicates the membrane interface separating the upper and lower chambers. Images were acquired after 24 h incubation. DAPI used to stain nuclei. **B.** Quantification of PDGF-BB-stimulated human pericyte migration following 24 h exposure to angiotensin II (AngII; 250 nM) and/or cigarette smoke extract (CSE; 10% concentration), expressed as percentage of total nuclei per well. **C.** Combined exposure of AngII and CSE inhibits pericyte proliferation in a dose-dependent manner. Proliferation assay quantified following 48 h exposure to AngII with or without CSE. * *P* < 0.05, ** *P* < 0.01 compared to the 10% CSE group.

## Discussion

Dysregulation of the RAS system, anchored on the angiotensin receptors, is implicated in a variety of common disorders such as cardiovascular disease, chronic renal dysfunction, and sepsis, yet the precise cellular context in which angiotensin receptors operate in the human lung has remained poorly defined. In this study, we integrate large-scale human transcriptomic analyses, *in vivo* mouse models of emphysema, and *in vitro* functional assays using primary human pericytes to establish a comprehensive, cell-resolved framework for angiotensin receptor biology in the lung. Leveraging publicly available single-nucleus and bulk RNA-sequencing datasets, we identify mutually exclusive and compartment-specific localization of the two predominant angiotensin receptors, with *AGTR1* almost exclusively expressed in lung pericytes and *AGTR2* restricted to a small epithelial subpopulation. Functional interrogation in experimental emphysema models and human pericyte co-cultures demonstrates that *AGTR1* signaling is a key regulator of pericyte abundance and behavior within the airspace microvasculature. Together, these computational, *in vivo*, and *in vitro* approaches converge to redefine angiotensin signaling in the lung as a pericyte-centered pathway with direct relevance to emphysema pathogenesis and therapeutic targeting.

Few prior studies have examined angiotensin receptor expression across lung cell types using validated reagents or modern transcriptomic platforms, contributing to uncertainty regarding receptor localization. Although angiotensin receptor-positive cells represent a small fraction of total lung cells, we show that they account for approximately one quarter of stromal cells across lung compartments. Within this population, *AGTR1* expression is concentrated in the perivascular niche, present in more than half of lung pericytes, whereas *AGTR2* expression is primarily expressed in alveolar epithelial type 2 cells. Earlier rodent studies identified bulk lung expression of the human equivalent *Agtr1a* isoform^41–43^ but lacked cellular resolution, while human *in situ* hybridization COPD studies suggested stromal and vascular smooth muscle expression of *AGTR1*^44^. By combining single-nucleus transcriptomics, pericyte isolation, and RNAscope validation, our study provides substantially greater anatomic specificity, distinguishing pericytes from other stromal populations and defining angiotensin receptor expression within the airspace compartment. These findings resolve longstanding ambiguity surrounding lung angiotensin receptor localization and provide a necessary foundation for interpreting prior pharmacologic studies.

The observation that *AGTR1* is preferentially expressed in pericytes rather than vascular smooth muscle cells represents a conceptual shift in how angiotensin signaling should be viewed in the lung. Pericytes are evolutionarily conserved perivascular cells that maintain intimate physical and functional relationships with the microvasculature, yet they remain incompletely characterized due to limited cell-specific markers and context-dependent phenotypes^45^. Prior lung pericyte research has largely focused on vascular development or pulmonary vascular disease^46–48^, with emerging work implicating these cells in fibrosis and asthma^49–51^. However, despite the central role of the alveolar microvasculature in emphysema pathogenesis, the contribution of pericytes to COPD has been largely unexplored. Our data here demonstrate for the first time that chronic cigarette smoke-induced is associated with reduced pericyte abundance in the airspace compartment, consistent with known microvascular loss and endothelial dysfunction in clinical and preclinical emphysema^52–54^.

Beyond a plausible function as a pericyte marker, we establish in preclinical platforms a direct functional consequence of AGTR1 modulation in the pericyte compartment. Restoration of pericyte-endothelial coupling by AGTR1 antagonism in cigarette smoke-exposed mice suggests that alveolar resiliency depends on the quantitative and functional integrity of pericyte-endothelial interactions, analogous to the strict pericyte:endothelial ratios required to maintain microvascular barrier function in the central nervous system^55^. While pericyte loss may both contribute to and result from endothelial dysfunction, our findings position pericytes as active participants in microvascular failure during emphysema progression. Both the migration of pericytes to endothelial cells and pericyte proliferation are attenuated by either CSE exposure or AGTR1 activation with some evidence of co-exposure synergism. In combination, these findings invoke a stabilization of the airspace microvascular pericyte-endothelial bed in the setting of CS injury that is conveyed by inhibition of AGTR1 signaling.

Using large public transcriptomic repositories, we were able to interrogate lung pericyte heterogeneity at scale. Unbiased subclustering identified multiple transcriptional states, with *AGTR1* expression detected across all pericyte subclusters, demonstrating that *AGTR1* is not restricted to a discrete pericyte subtype. Quantitative differences in *AGTR1* expression were observed among subclusters, with modest enrichment in a single cluster, but broad expression across transcriptional states. In contrast, expression of the canonical contractile marker *ACTA2* also varied across pericyte subclusters, and patterns of *ACTA2* expression did not align with *AGTR1* enrichment. Together, these findings indicate that *AGTR1* expression spans multiple pericyte transcriptional states and is not coupled to classical contractile identity, supporting its role as a broadly relevant signaling receptor within the lung pericyte population.

Although *AGTR1* expression is most highly enriched in lung pericytes, we also detected lower-level expression in multiple fibroblast populations, with disease-associated dysregulation that varied by stromal compartment. Across the HLCA dataset, *AGTR1* dysregulation was most pronounced in peribronchial fibroblasts, which exhibited significantly higher expression in chronic rhinitis, interstitial lung disease, and pulmonary fibrosis compared with normal lungs, with consistent effect sizes across contrasts. In contrast, pericytes, alveolar fibroblasts, adventitial fibroblasts, and vascular smooth muscle cells showed only modest or trend-level disease-associated changes in *AGTR1* expression, with no post hoc comparisons surviving multiple-testing correction. Importantly, the presence of disease-associated *AGTR1* dysregulation in select fibroblast populations does not contradict a pericyte-centered model of lung angiotensin signaling, but rather suggests that receptor localization and disease responsiveness may be uncoupled across stromal compartments.

Targeted analysis of an independent single-nucleus lung dataset enriched for COPD and IPF further supported disease-associated engagement of *AGTR1* within mesenchymal compartments, revealing increased *AGTR1* expression in fibroblasts from COPD patients and a higher prevalence of *AGTR1*⁺ cells across IPF donor lungs. While mean *AGTR1* expression within individual cell types did not differ significantly between IPF and control donors, the broader distribution of *AGTR1*⁺ cells in IPF suggests widespread, albeit heterogeneous, activation of this pathway. Subclustering of pericytes in both datasets revealed no robust disease-specific dysregulation after correction for multiple testing, although select subclusters exhibited nominal trends, highlighting potential subtype-specific sensitivity that may require larger cohorts to resolve. Collectively, these findings indicate that disease-associated *AGTR1* dysregulation is not uniform across the stromal compartment, but instead is concentrated in specific fibroblast populations and variably engaged across pericyte states, consistent with context-dependent roles for *AGTR1* signaling in lung injury and repair.

Functional studies provide mechanistic support for these observations. Both angiotensin II and cigarette smoke exposure impair pericyte survival and migration toward endothelial cells, indicating convergent inhibitory effects on pericyte behaviors essential for vascular attachment. The synergistic suppression of pericyte proliferation by combined angiotensin II and cigarette smoke exposure suggests that chronic inflammatory and neurohumoral signaling may cooperatively destabilize the microvasculature in COPD. While proinflammatory effects of angiotensin II are well recognized, stromal cell contributions are less frequently considered.

Given the inflammatory milieu of COPD, it will be important to determine whether *AGTR1*-mediated effects extend to immune-stromal crosstalk. In parallel, the identification of *AGTR2* expression in a discrete alveolar epithelial subpopulation raises the possibility that AGTR1 blockade may simultaneously relieve pericyte dysfunction while permitting *AGTR2*-mediated epithelial repair, consistent with our prior observations in acute lung injury^38^.

Several limitations warrant consideration. Transcriptomic analyses depend on RNA quality and dataset composition, and disease-specific cell populations remain underrepresented in some cohorts, limiting statistical power for detecting cell type- and subtype-specific disease effects. Nonetheless, the consistency of angiotensin receptor localization across independent datasets strengthens our conclusions. In addition, the more pronounced reduction in pericyte abundance relative to endothelial cells following chronic cigarette smoke exposure raises the possibility that these cell types differ in their capacity to engage protective or adaptive responses during prolonged injury. Defining the molecular basis of this differential vulnerability will be an important focus of future studies. Finally, future investigations incorporating larger disease-focused cohorts and spatial transcriptomic approaches will be important to refine compartment-specific signaling dynamics and to resolve context-dependent cellular interactions within the lung microvasculature.

In summary, by integrating computational analysis of large human transcriptomic datasets with mechanistic *in vivo* and *in vitro* experimentation, we redefine angiotensin receptor biology in the lung. Our findings establish *AGTR1* as a selective and functionally relevant marker of lung pericytes, demonstrate that pericyte loss and endothelial uncoupling are features of emphysema, and show that AGTR1 antagonism restores key aspects of microvascular integrity. These results provide a unifying cellular framework for interpreting prior ARB studies in lung disease and suggest that therapeutic efficacy may critically depend on preserving or restoring pericyte-endothelial interactions within the airspace microvasculature. More broadly, this work highlights the power of public transcriptomic resources to uncover cell-specific signaling pathways and illuminate the roles of anatomically and functionally elusive populations such as pericytes in complex lung disorders.

## Methods

### Sex as a biological variable

Sex was explicitly considered as a biological variable throughout the study design and analysis. Human transcriptomic analyses incorporated samples from both male and female donors, and sex was included as a covariate in all donor-level statistical models evaluating *AGTR1* expression, disease associations, and aging effects, thereby controlling for potential sex-related confounding. Experimental mouse studies included animals of the same sex within each experiment to reduce biological variability and increase statistical power; this approach was selected based on established precedents for emphysema and cigarette smoke-exposure models, in which sex-specific differences in baseline lung structure and injury responses can complicate interpretation. While the use of a single sex in these experiments limits direct assessment of sex-dependent effects, the concordance between human multi-sex data and murine findings supports the broader relevance of the conclusions. Collectively, these design choices ensure that sex-related influences were either directly modeled or experimentally controlled, and the findings are expected to be broadly applicable across sexes, while motivating future studies explicitly powered to assess sex-specific mechanisms.

### Human lung data acquisition and study populations

We obtained HLCA v2^31^, LungMAP^32^, GTEx v8^37^, and IPF/COPD^39,40^ single-cell datasets from public repositories (CellxGene, LungMAP, GTEx Portal, GEO GSE136831), with exact accession links provided under *Data Availability*. Donors represented a diverse range of ancestries (predominantly European), ages (31 weeks to 80 years), and clinical backgrounds (**Table S4**).

### Single-nucleus RNA-sequencing processing and quality control

Single-cell and single-nucleus RNA-seq data were processed using *SingleCellExperiment* (version 1.24.0) in R (version 4.3). Counts were normalized using logNormCounts from *scuttle*^56,57^ (version 1.12.0) and sequencing-depth factors were computed using *scran*. Per-cell and per-feature QC metrics (mitochondrial percentage, library size) were added using *scuttle*.

QC filtering thresholds were dataset-specific:

- **HLCA v2:** Mitochondrial percentages were uniformly low; cells were filtered using perCellQCFilters without mitochondrial-specific thresholds.
- **LungMAP 10x:** Removed cells with >25% mitochondrial reads or <1,000 UMIs.
- **IPF/COPD (GSE136831):** Removed cells with >25% mitochondrial reads and <1,000 UMIs.

Only post-QC cells were included in angiotensin receptor analyses.

### Angiotensin receptor localization analysis

#### Primary analysis in HLCA and enrichment

*AGTR1* and *AGTR2* expression was evaluated across HLCA annotation levels (compartment and cell types). To assess differential localization of angiotensin receptors, we fit donor-level linear models of normalized expression across annotation groups and corrected for multiple testing (FDR). For enrichment analysis, we applied a two-sided Fisher’s exact tests on angiotensin receptor expression across compartments and cell types, with FDR correction applied within each gene-annotation combination.

#### LungMAP replication via SCANVI label transfer

To harmonize LungMAP (query) with HLCA (reference), we employed SCANVI^58^ (scvi-tools^59–61^; version 1.3.3). Log-normalized HLCA reference data were subset to 2,000 highly variable genes. Log-normalized LungMAP cells were projected into the shared latent space, and labels were transferred based on a posterior probability ≥ 0.90.

### Pericyte-specific subclustering and proximity analysis

#### Pericyte subsetting and batch correction

Pericytes (n=11,680 cells; 194 donors) were subset from the HLCA v2 full dataset based on the finest-level annotations provided by the atlas. To account for technical and biological variation across the integrated studies, batch correction was performed using Harmony (harmonypy ^62^; v0.0.10). The correction model incorporated donor, study source, sequencing platform, sex, age, and self-reported race as covariates. The resulting Harmony-corrected latent space was used for all downstream pericyte subclustering and transcriptomic proximity calculations.

#### Perivascular reference subset for airspace proximity scoring

To quantify the transcriptomic proximity of pericytes to the alveolar-capillary interface, we generated a specialized “perivascular” reference subset from the HLCA v2 full dataset. This subset provided the compartmental reference populations necessary for computing latent space centroids. In addition to pericytes, this reference included:

- **Alveolar Epithelium:** AT1 and AT2 cells.
- **Vascular Endothelium:** Capillary aerocytes, general capillaries, arterial, and venous endothelial cells.
- **Lymphatic Endothelium:** Differentiating, mature, and proliferating lymphatics.
- **Vascular Smooth Muscle (vSMC):** General smooth muscle, smooth muscle, and activated stress-response smooth muscle.

#### Airspace proximity scoring

To quantify pericyte proximity to the alveolar airspace, we calculated an *airspace proximity score (*𝑆_𝑖_) based on the mean cosine similarity (**Equation 1**) between pericyte Harmony-embeddings (𝑧_𝑖_) and the centroids of reference airspace classes (𝒞 = {AT1, AT2, EC_aerocyte_,EC_general_}):

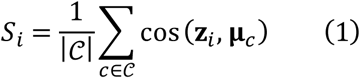

Airspace proximity was compared between *AGTR1*⁺ and *AGTR1*-negative pericytes using donor-level aggregation. Linear regression models were fit with donor age and sex as covariates, and results corrected for multiple testing (FDR).

#### Technical dropout and denoising

To determine whether the observed frequency of *AGTR1*-negative pericytes exceeded expectations due to technical dropout, we implemented a matched-gene dropout expectation model. Genes with similar mean expression and coefficient of variation to *AGTR1* were selected as controls (**Equation 2**). For each pericyte 𝑖, the expected probability of zero detection (*p*_𝑖_) was estimated as the fraction of zeros across matched-gene set (𝑀):

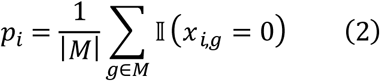

The observed number of *AGTR1* zeros across all pericytes was compared against a Poisson-binomial null distribution parameterized by *p*_𝑖_. Enrichment (excess sparsity) was assessed using a one-sided exact test implemented via fast Fourier transform.

To further evaluate the impact of technical under calling, *AGTR1* expression was denoised using a negative binomial variational autoencoder (scVI^61^). Models were trained on pericytes alone or an expanded perivascular compartment (including endothelial and smooth muscle cells), with “study” as a batch covariate. Denoised *AGTR1* estimates were calibrated to preserve the observed population-level prevalence to ensure biological realism. Associations between denoised *AGTR1* status and airspace proximity were then evaluated using donor-clustered linear models to account for inter-individual variability.

#### De novo pericyte subclustering

Neighborhood graphs were constructed using cosine distance in the Harmony -corrected latent space. To identify transcriptionally distinct pericyte states, we performed Leiden community detection (implemented via Scanpy^63^; version 1.11.4), with the clustering resolution fixed at 0.6 to optimize the balance between biological interpretability and cluster granularity.

Differentially expressed marker genes for each subcluster were identified using the Wilcoxon rank-sum test (scanpy.tl.rank_genes_groups) with Benjamini-Hochberg correction for multiple testing. Only genes with a log-fold change > 0.25 and an adjusted *p*-value < 0.05 were considered for marker-based characterization.

### Age-associated variation in AGTR1 expression

#### HLCA single-nucleus analysis

To assess cell-type-specific age variation, we analyzed transcriptomic data from the HLCA v2. To account for the hierarchical structure of the data and avoid pseudo replication, *AGTR1* expression was summarized at the donor–cell type level. For each donor-cell type pair, we calculated mean expression (average log-normalized *AGTR1* levels) and cell fraction (proportion of *AGTR1*⁺ cells). Estimation was restricted to donor-cell type combinations meeting minimum cell (n ≥ 20) and donor count (n ≥ 3) thresholds to ensure statistical robustness.

Associations between donor age and *AGTR1* expression were first evaluated using Spearman correlation with Benjamini-Hochberg FDR correction across cell types. To assess non-linear age effects, generalized additive models and nested linear models comparing linear versus spline-based age terms were fitted with adjustment for sex and self-reported ethnicity, with significance assessed by likelihood ratio or smooth-term tests and corrected for multiple testing.

#### Cell-type deconvolution of bulk tissue

To extend these findings to the GTEx bulk lung dataset, deconvolution was performed using BayesPrism^64^ (version 2.2.2) with HLCA v2 as the single-cell reference (**Figure S5**). Marker genes were selected using get_mean_ratio2 from DeconvoBuddies^65^. To ensure high-quality estimates, we excluded ribosomal, mitochondrial, and X/Y chromosome genes, as well as the non-coding RNA *MATAL1* following BayesPrism guidelines. The new.prism function was run with an outlier cutoff of 1% and an outlier fraction of 10%.

#### Correlation in bulk lung tissue

For the 578 GTEx lung samples, we evaluated the relationship between receptor expression and age using linear models. Models were fit for log_10_-transformed TPM expression of each angiotensin receptor as a function of donor age. *P*-values were Bonferroni-corrected to maintain a stringent threshold for significance across angiotensin receptors.

### Disease association in human lung

#### Cell type and pericyte subcluster disease association in HLCA

For stromal cell types, donor-level mean *AGTR1* expression and the fraction of *AGTR1*⁺ cells were calculated from log-normalized single-cell data, restricting analyses to donors aged ≥20 years, excluding cancer diagnoses, and requiring ≥10 cells per donor–cell-type combination and ≥3 donors per group. Disease-associated differences were evaluated using Kruskal–Wallis tests followed by Dunn’s post hoc comparisons with FDR correction, with effect sizes estimated from test statistics. To assess whether transcriptional heterogeneity masked disease-associated *AGTR1* patterns within pericytes, pericytes were further stratified into harmonized transcriptional subclusters using HLCA-derived Leiden annotations, and donor-level *AGTR1* metrics were computed for each subcluster under the same filtering criteria. Disease associations across pericyte subclusters were assessed using the same nonparametric framework, with all analyses conducted at the donor level to account for inter-individual variability.

#### IPF and COPD angiotensin receptor expression analysis

A previously published human lung single-cell RNA-seq dataset comprising control, COPD, and IPF donor lungs (GSE136831) was reprocessed and normalized. Donor-level expression of *AGTR1* and *AGTR2* was assessed across cell types and disease states using mean normalized expression, with group differences evaluated by two-sided *t* tests or one-way ANOVA as appropriate, and donor-level *AGTR1* positivity defined using a minimal expression threshold.

Pericytes were subset from this dataset, batch-corrected at the donor level using Harmony, and transcriptionally aligned to HLCA pericyte subclusters using a scANVI-based reference-query framework to obtain probabilistic subcluster assignments and latent embeddings. Donor-level *AGTR1* expression and *AGTR1*⁺ cell fractions were then computed for each predicted subcluster, and disease-associated differences were assessed using Kruskal–Wallis tests with Dunn’s post hoc comparisons and FDR correction, restricting analyses to adequately represented subclusters.

### Cigarette smoke exposure

Six- to eight-week-old male AKR/J mice were assigned to filtered-air or cigarette-smoke exposure groups. Smoke exposure used 2R4F reference cigarettes (University of Kentucky) burned for two hours per day, five days per week, for 6-7 weeks using a Teague TE-10 smoking machine, maintaining 90 mg/m³ total suspended particulates and 350 ppm CO.

### Mouse pericyte isolation

Protocol was adapted from one graciously provided by Carole L Wilson, PhD at the University of Wisconsin-Madison^66^. Briefly, 3-month-old C57/BL6 mice from Jax (Strain #: 000664) were anaesthetized with ketamine/xylazine (concentrations) before cervical dislocation. The chest cavity was opened, and the portal vein was severed to allow easy flushing of the lungs. The ribcage was dissected to expose the heart, lungs, and trachea. The trachea was sutured shut, and the lungs were perfused with sterile PBS via the right ventricle. Once sufficient clearance of blood was assessed via lightening of tissue color, lungs were removed and placed in DMEM. Once harvesting was finished, lung lobes were removed from the trachea and bronchi and sliced into small (1-2 mm) pieces using a scalpel. Tissue pieces were placed in media containing 0.5 mg/mL Liberase TL (Roche, #05-401-020-001) and incubated at 37°C for 2 hours (or until sufficiently digested) on a rotary shaker. Cell suspension was filtered once with a 100 μm filter and subsequently with a 70 μm filter to attain single cell suspensions. Single cell suspension was centrifuged and resuspended in ACK red blood cell lysis buffer to remove residual erythrocytes. Cells were washed twice with HBSS and then resuspended in MACS incubation buffer (MACS BSA Stock solution [Miltenyi Biotec #130-091-376] diluted 1:20 in MACS wash buffer [Miltenyi Biotec #130-091-222]). PE-conjugated anti-mouse PDGFRβ (CD140b) microbeads (Miltenyi Biotec #130-096-267) were added (1:5) to cell suspension and incubated at 4°C for 10 minutes before washing and resuspension in MACS wash buffer.

Magnetic anti-PE microbeads were added to cell suspension (1:5) and incubated at 4°C for 15 minutes. Magnetic separation was performed using a QuadroMACS magnet (Miltenyi Biotec #130-090-976) and LS columns (Miltenyi Biotec #130-042-401). Flowthrough was collected as the PDGFRβ-negative population and placed in TRIzol (Invitrogen; #15596026) for RNA isolation. Column-bound cells were washed with MACS wash buffer before elution into TRIzol and subsequent RNA isolation.

### Quantitative real-time PCR analysis

Total RNA was isolated from mouse lung pericytes or whole lung tissue using TRIzol reagent according to the manufacturer’s instructions, followed by DNase treatment. First-strand cDNA was generated using Invitrogen reverse transcription kits (SuperScript III; #18080-044).

Quantitative real-time PCR was performed in technical triplicate using TaqMan probes (below; Applied Biosystems) on Applied Biosystems real-time PCR systems (StepOnePlus and ABI Fast 7500). Relative gene expression was calculated using 2^-ΔΔCt^ method, with GAPDH used as the reference gene. For pericyte analyses, the PDGFRβ-negative cell population was used as the reference sample.

| Gene ID | Probe ID |
| --- | --- |
| *Agtr1a* | Mm01957722 |
| *Pdgfrb* | Mm00435553 |
| *Vcan* | Mm01283063 |
| *Pecam* | Mm01242576 |
| *Des* | Mm00802455 |
| *Gapdh* | 4352339E |

### Lung ViewRNA in situ

Lung sections on SuperFrost Plus (+) slides were stained using the manufacturer’s protocol for tissue ViewRNA (ThermoFisher, QVT0700). Briefly, sections were deparaffinized before antigen retrieval in ViewRNA antigen retrieval buffer. After subsequent washes and protease pretreatment, slides were labeled using the following probe and dye combinations: <u>Mouse:</u> Agtr1a, VB6-12945, Alexa 647; Pdgfrb, VB4-3112504, Alexa 488; Higd1b, VB1-3040702, Alexa 546. Human: AGTR1, VA1-17758, Alexa 546; HIGD1B, VA6-3178101, Alexa 647. After washes and amplification steps, slides were then incubated in DAPI solution before addition of ProLong Diamond mounting media (Invitrogen, P36971) and placement of a coverslip (THORLABS, #1.5H) Slides were imaged on an Olympus FV3000RS confocal microscope with a 20x air objective (UPLSAPO20X).

### Lung tissue immunohistochemistry

Lung sections were mounted to SuperFrost Plus (+) slides. Sections were deparaffinized before antigen retrieval (Vector Labs, H-3300) for 15 minutes at ∼100°C. After antigen retrieval, slides were blocked in a 5% BSA/0.1% Triton X-100/PBS solution for 45 minutes at room temperature. Slides were then incubated in primary antibody solution in PBS/0.02% Triton X-100 at 4°C overnight. Antibody dilutions and information are included in the table below. After primary antibody incubation, slides were washed in PBS and incubated in secondary antibody solution made in PBS. Slides were then incubated in DAPI solution before addition of ProLong Diamond mounting media (Invitrogen, P36971) and placement of a coverslip (THORLABS, #1.5H). Slides were imaged on an Olympus FV3000RS confocal microscope with a 30x oil immersion objective (UPLSAPO30XS). Cells were counted using the Fiji CellCounter plugin.

| **Host species** | **Antibody target** | **Vendor, catalog number** | **RRID** | **Dilution** |
| --- | --- | --- | --- | --- |
| Goat | Mouse PDGFRβ | R&D Systems, AF1042 | AB_2162633 | 1:200 |
| Rat | Mouse PECAM | Synaptic Systems, HS-351117 | AB_3719883 | 1:250 |
| Rabbit | Mouse NG2 | Sigma Aldrich, ZRB-5320 | AB_3677668 | 1:75 |
| Goat | Human PDGFRβ | R&D Systems, AF385 | AB_355339 | 1:50 |
| Rabbit | Human NG2 | Abcam, AB139406 | AB_2847916 | 1:100 |
| Mouse | Human PECAM | Abcam, AB9498 | AB_307284 | 1:100 |
| Donkey | Goat IgG | ThermoFisher, A11057 | AB_2534104 | 1:500 |
| Donkey | Rabbit IgG | ThermoFisher, A32795 | AB_2762835 | 1:500 |
| Donkey | Rat IgG | ThermoFisher, A21208 | AB_2535794 | 1:500 |
| Donkey | Mouse IgG | ThermoFisher, A32766 | AB_2762823 | 1:500 |

### Pericyte transwell migration assay

Human pericytes were a gift from Dr. Vinicio de Jesus Perez and were isolated according to his previously publications^67^. Cells were cultured in Human Pericyte media (ScienCell, Cat#: 1201) at 37°C in 5% CO2 before plating for experiments. Transwell membranes (Falcon, #353097) were coated in gelatin (Cell Biologics, #6950) before assay. Cells were plated 20000/well and allowed to adhere overnight. The next day, media in the upper well was changed to serum free human pericyte media, and lower chamber media was changed to complete media containing PDGF-BB (Sigma P3201, 50 ng/mL) and ANGII (Sigma A9525, 500nM) or cigarette smoke extract. Membranes were incubated a further 24 hours to allow for cell migration before fixation in 4% PFA in PBS and incubation in DAPI solution. Membranes were excised from plastic supports and placed on a slide with ProLong Diamond mounting media (Invitrogen, P36971).

Images were collected on an Olympus FV3000RS confocal microscope with a 10x air objective (UPLSAPO10X2) through the span of the membrane to capture cells on both sides. To obtain cell counts and migration percentages, Z-stacks were generated and analyzed with Fiji/ImageJ.

### Pericyte proliferation assay

Human pericytes were cultured in Pericyte medium (ScienCell, Cat#: 1201) up to passage 8. Cells were plated in 96 well plate with 5000 cells plated per well. The next day, cell medium was changed to serum free media and incubated overnight. After overnight incubation in serum free media, Ang II was added to wells and after 30 min incubation. Cigarette smoke extract was then added and cells were incubated for 24 hours before fluorescence was read. Cell proliferation was assessed using the CyQUANT NF cell proliferation assay from Molecular Probes (C35006).

### Graphics

All plots were generated in R (version 4.3) or Python (version 3.10) unless otherwise stated. Venn diagrams comparing overlap between *AGTR1* and *AGTR2* expression were generated in R using ggvenn^68^ (version 0.1.10). Dot plots depicting average expression and the proportion of angiotensin receptor–positive cells were produced using the Seurat^69^ DotPlot function (version 5.0.1). Line graphs, scatter plots, box plots, and bar plots were generated in R using ggpubr^70^ (version 0.6.0). Heatmap-style enrichment plots were created using ggplot2^71^ (version 3.5.0) with ggfittext (version 0.10.2) and ggpubr for annotation and layout. Additional exploratory visualizations were generated in Python using seaborn^72^ (version 0.13.2). Molecular analysis results were additionally visualized using GraphPad Prism (version 9.0) and FIJI/ImageJ for quantitative plotting and image-based analysis, respectively.

### Statistics

All statistical analyses were performed in R (version 4.3) and Python (version 3.10). Unless otherwise specified, statistical tests were two-sided, and multiple-testing correction was applied using Benjamini-Hochberg FDR correction. A corrected *p* value < 0.05 was considered statistically significant.

For single-nucleus RNA-seq analyses, inference was conducted at the donor level to avoid pseudo replication. Donor-level mean normalized *AGTR1* expression and the fraction of *AGTR1*⁺ cells were calculated for each cell type or pericyte subcluster, subject to minimum cell and donor count thresholds as specified.

Comparisons of *AGTR1* expression across cell types, compartments, disease states, or pericyte subclusters were performed using Kruskal-Wallis tests, followed by Dunn’s post hoc tests with FDR correction. Effect sizes for nonparametric tests were estimated from standardized test statistics. For enrichment analyses of receptor expression, two-sided Fisher’s exact tests were applied with FDR correction within each gene-annotation combination.

Associations between *AGTR1* expression and continuous variables (e.g., age, airspace proximity score) were assessed using linear regression models or Spearman rank correlations, as appropriate. Multivariable models included adjustment for relevant covariates (e.g., age, sex, self-reported ethnicity), and donor-clustered models were used where indicated to account for inter-individual variability. For non-linear age effects, generalized additive models and nested linear models comparing linear versus spline-based age terms were fitted, with significance assessed using likelihood ratio tests or smooth-term tests and corrected for multiple testing.

To assess whether the frequency of *AGTR1*-negative pericytes exceeded expectations from technical dropout, the observed number of zero counts was compared against a Poisson-binomial null distribution parameterized by per-cell dropout probabilities derived from matched control genes. One-sided exact *p* values were computed using fast Fourier transform-based methods. Associations involving denoised *AGTR1* expression derived from scVI were evaluated using donor-clustered linear models.

For bulk RNA-seq analyses of GTEx lung tissue, associations between receptor expression and age were evaluated using linear models fit to log-transformed TPM values. Bonferroni correction was applied to account for testing across angiotensin receptor genes.

For mouse and whole-cells studies, comparisons between two groups were performed using unpaired, two-tailed *t* tests. Comparisons among more than two groups were conducted using one-way analysis of variance (ANOVA), while comparisons involving two or more independent variables were performed using two-way ANOVA in the GraphPad Prism software. Statistical tests were performed on biological replicates, and *P* < 0.05 were considered statistically significant.

Sample sizes, exact statistical tests, and adjusted *p* values are reported in figure legends and corresponding tables.

### Study approval

Adult AKR/J mice (Jackson Laboratory) were housed in an AAALAC-accredited facility. The Johns Hopkins University School of Medicine Institutional Animal Care and Use Committee approved animal procedures. All human data were drawn from publicly available, consented studies as described in the original publications^31,32,37,39,40^.

## Data availability

The HLCA is a publicly available resource located at https://hlca.ds.czbiohub.org/. We downloaded HDF5 files associated with the HLCA version from CellxGene https://cellxgene.cziscience.com/collections/6f6d381a-7701-4781-935c-db10d30de293. The LungMAP is a publicly available resource located at https://lungmap.net/. We downloaded data from Wang et al.^32^ at https://data-browser.lungmap.net/explore/projects/20037472-ea1d-4ddb-9cd3-56a11a6f0f76. For COPD and IPF data, we downloaded raw counts and metadata from GEO (GSE136831)^39,40^.

Computational analyses were performed using NSF-supported ACCESS resources^73^ on the Bridges-2 system^74^. All code, jupyter-notebooks, and results are available through GitHub at https://github.com/heart-gen/angiotensinII_lung.

## Declarations

### Author contributions

Conceptualization, KJMB and ERN; Methodology, KJMB, EG, MS, SG, AM, and ERN; Software, KJMB; Formal Analysis, KJMB, EG, and AM; Data curation, KJMB, MS, and SG; Writing – original draft preparation, KJMB and ERN; Writing – review and editing, KJMB, EG, MS, and ERN; Visualization, KJMB and EG; Supervision, ERN; Project administration, KJMB; Funding acquisition, KJMB and ERN.

## Acknowledgments

We thank Vinicio Jesus De Perez for assistance with the optimization of human pericyte studies. The results presented here are based in part on data generated by the LungMAP Consortium [U01HL122642] and obtained from the LungMAP Data Portal (www.lungmap.net) on February 2nd, 2022. The LungMAP Consortium and the LungMAP Data Coordinating Center (1U01HL122638) are funded by the NHLBI.

This work used the Bridges-2 system at the Pittsburgh Supercomputing Center through allocation BIO250079 from the Advanced Cyberinfrastructure Coordination Ecosystem: Services & Support (ACCESS) program, which is supported by National Science Foundation grants #2138259, #2138286, #2138307, #2137603, and #2138296.

