## Supplementary Information for "Convergence of Angiotensin Signaling on Lung Pericyte and Stromal Behaviors"

### **Supplementary Notes**

### **Supplementary Figure**

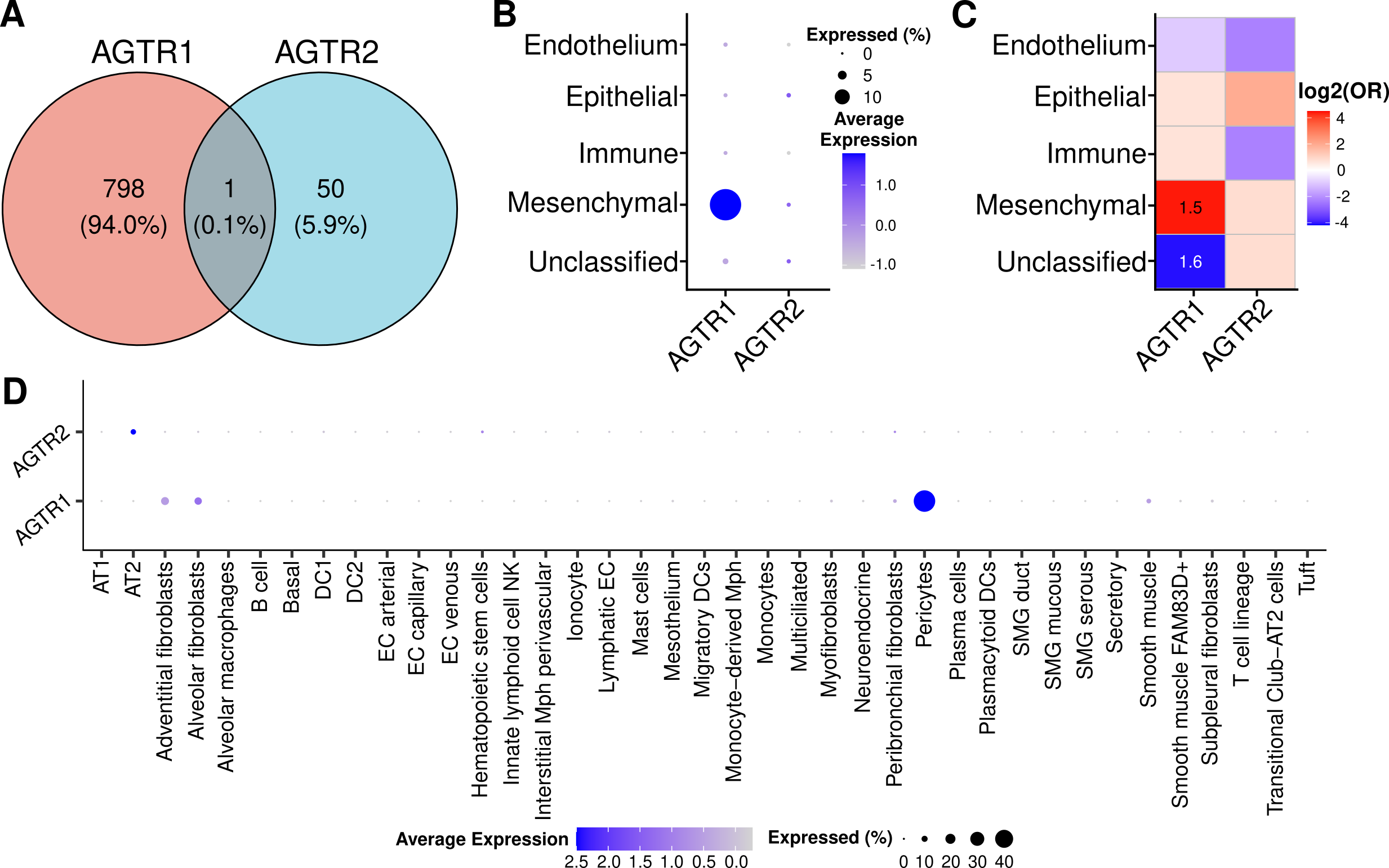

**Figure S1: Replication of AGTR1 expression for mesenchymal/stromal compartment- and pericytes in the human lung.** **A.** Venn diagram showing rare occurrences of AGTR1 and AGTR2 co-expression (n=9 individuals). **B.** Dot plot showing the percentage and average expression of AGTR1- and AGTR2-positive cells across lung compartments (n=9). **C.** Heatmap showing significant enrichment (two-sided, Fisher’s exact test) of AGTR1- and AGTR2-positive cells across lung compartments (n=9). The color intensity of enrichment heatmaps represents log2 of odds ratio (OR) with red indicating enrichment and blue indicating depletion. Significantly enrichment compartments are annotated with -log10(false discovery rate). **D.** Dotplot showing the percentage and average expression of AGTR1- and AGTR2-positive cells across lung cell types (n=9).

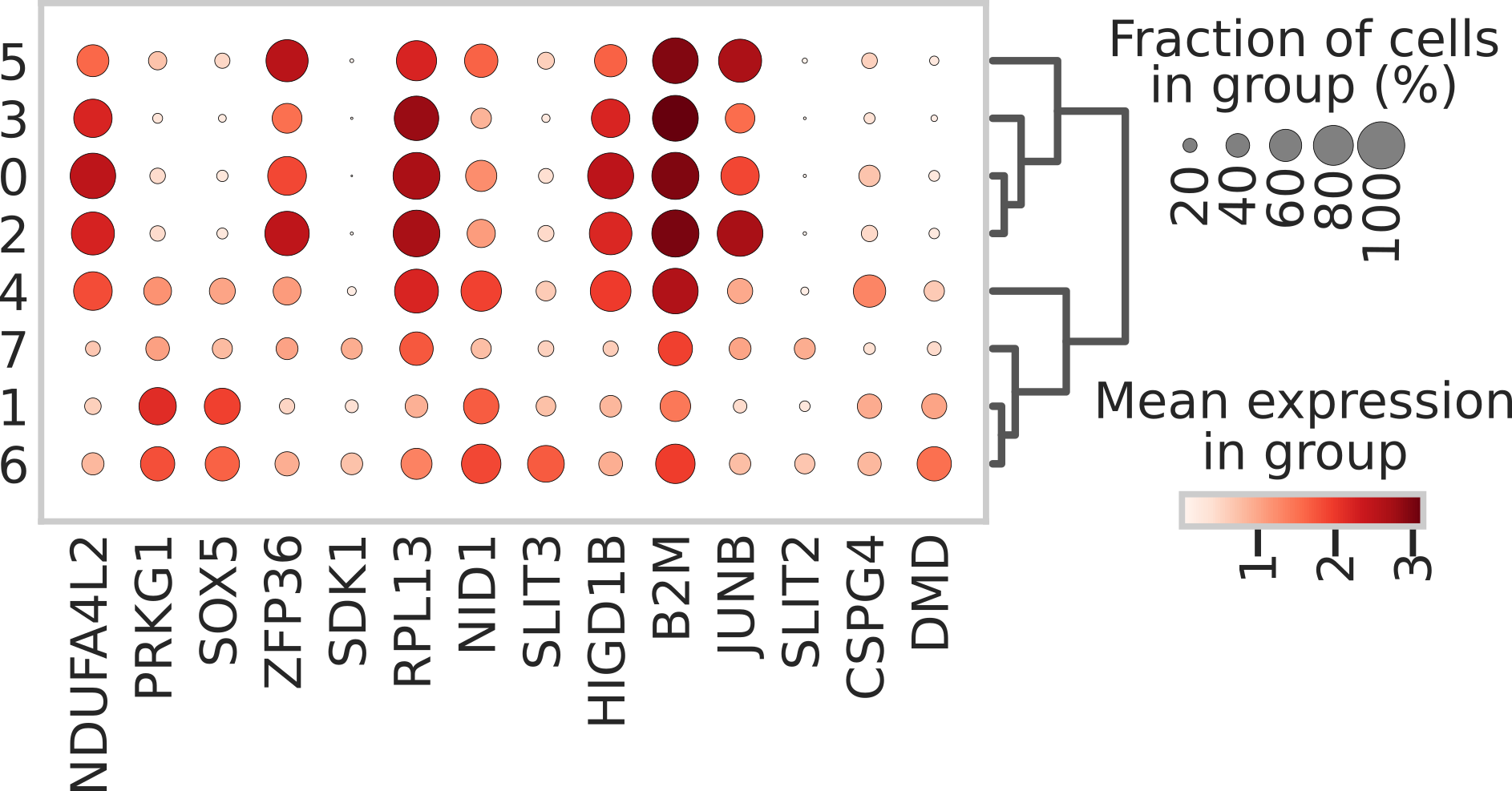

**Figure S2: Pericyte subclustering marker genes.** Dot plot showing the top two marker genes per Leiden subcluster (clusters 0-7) identified via Wilcoxon rank-sum test. Dot size represents the percentage of cells expressing the gene, and color intensity indicates the mean expression level within the cluster.

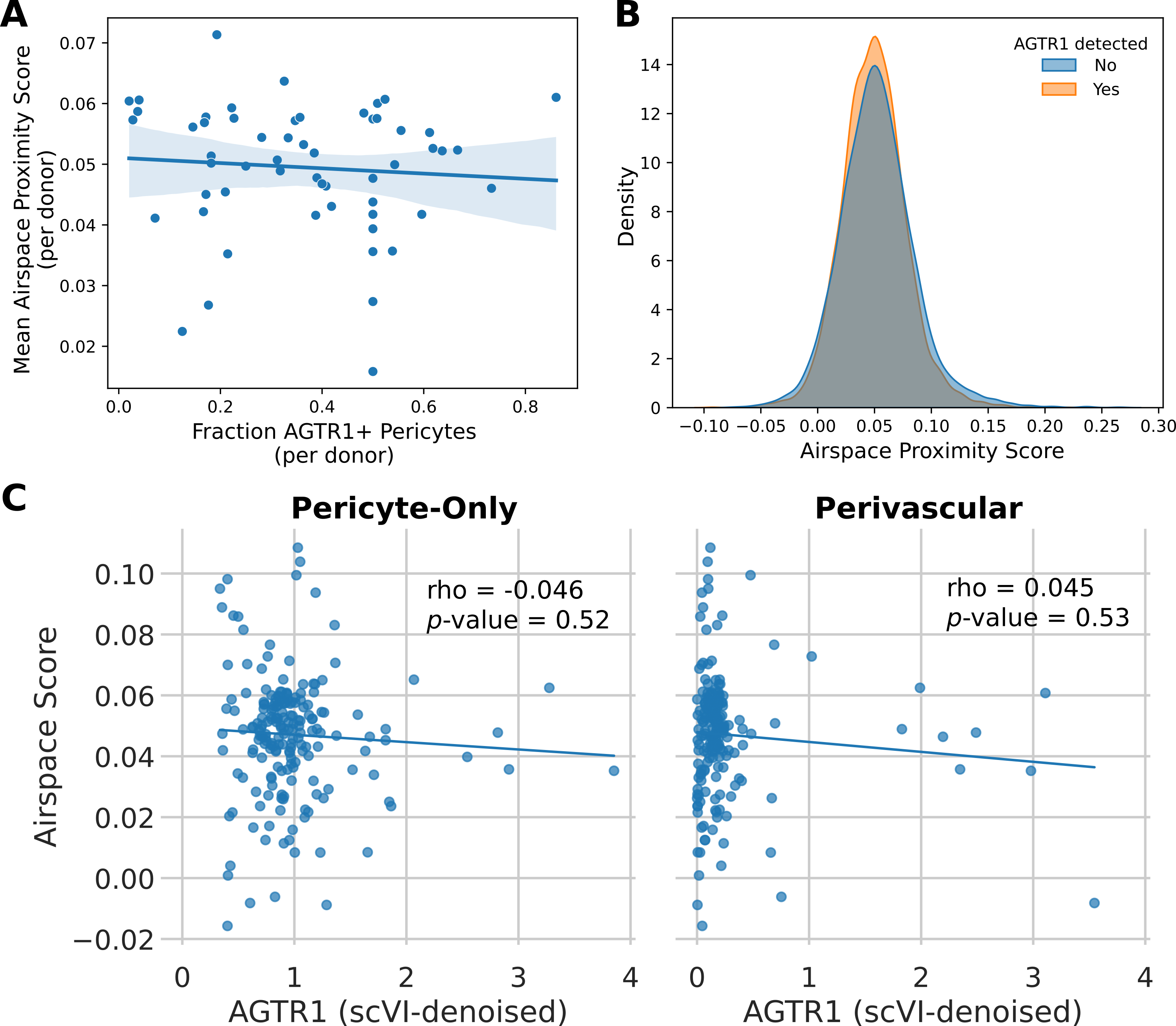

**Figure S3: No evidence for compartmental or spatial segregation of AGTR1-negative pericytes.** **A.** Scatter plot showing no correlation with airspace affinity and fraction of AGTR1-positive cells. **B.** Ridge plot showing overlap of airspace affinity for AGTR1+ and AGTR1-negative cells. **C.** Scatter plot showing no correlation with imputed AGTR1 and airspace affinity using pericyte- and perivascular-trained models.

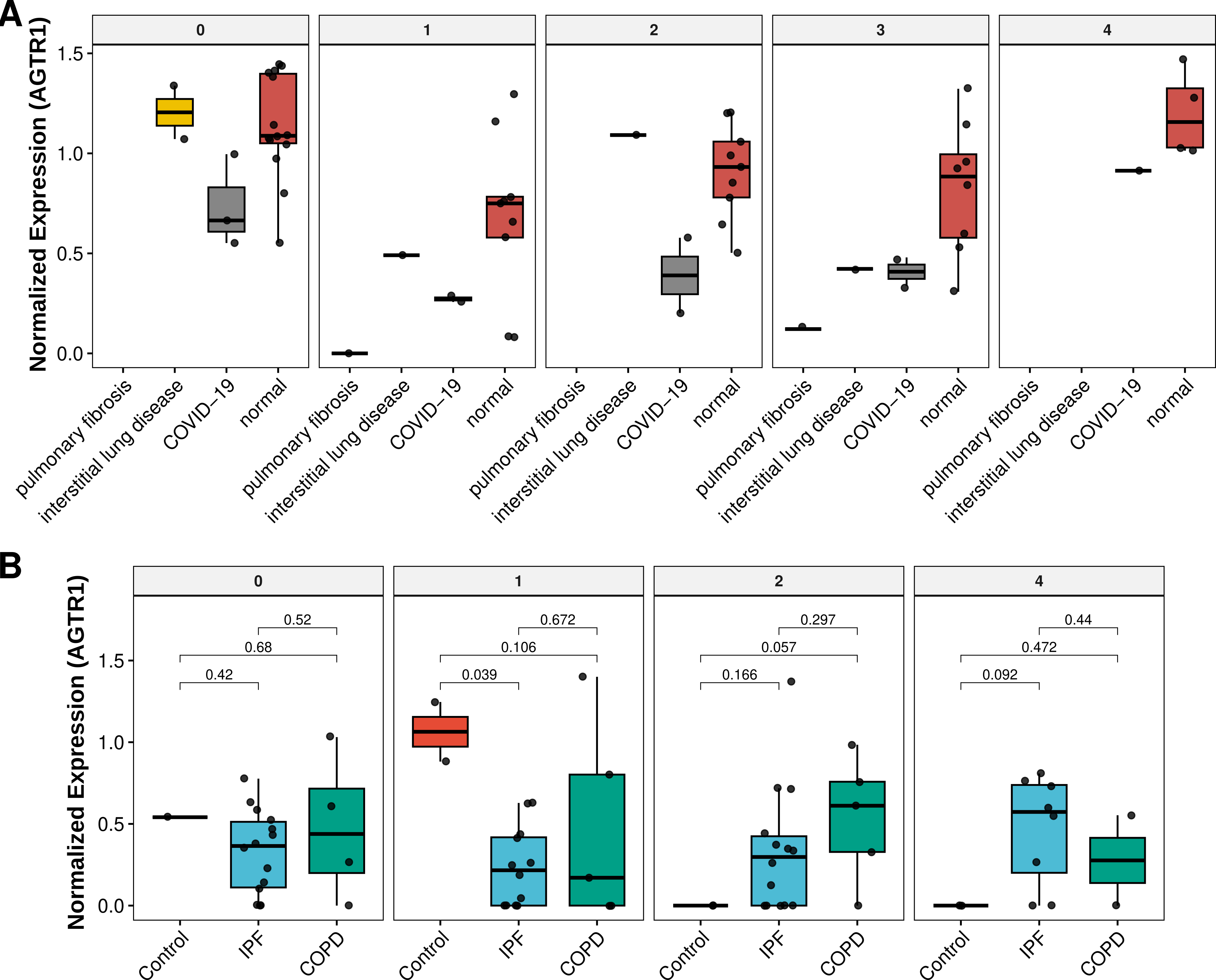
**Figure S4: Analysis of AGTR1 expression across pericyte subclusters.** **A.** Boxplots of AGTR1 expression in pericyte subclusters across disease states in the HLCA dataset. **B.** Boxplots of AGTR1 expression across transcriptionally defined pericyte subclusters, highlighting heterogeneity within the pericyte compartment in an independent single-nucleus lung dataset, separated by diagnosis (control, COPD, or IPF). Data are shown as log-normalized expression values. Boxplots display median, interquartile range (IQR), and 1.5× IQR whiskers; individual points represent donor means.

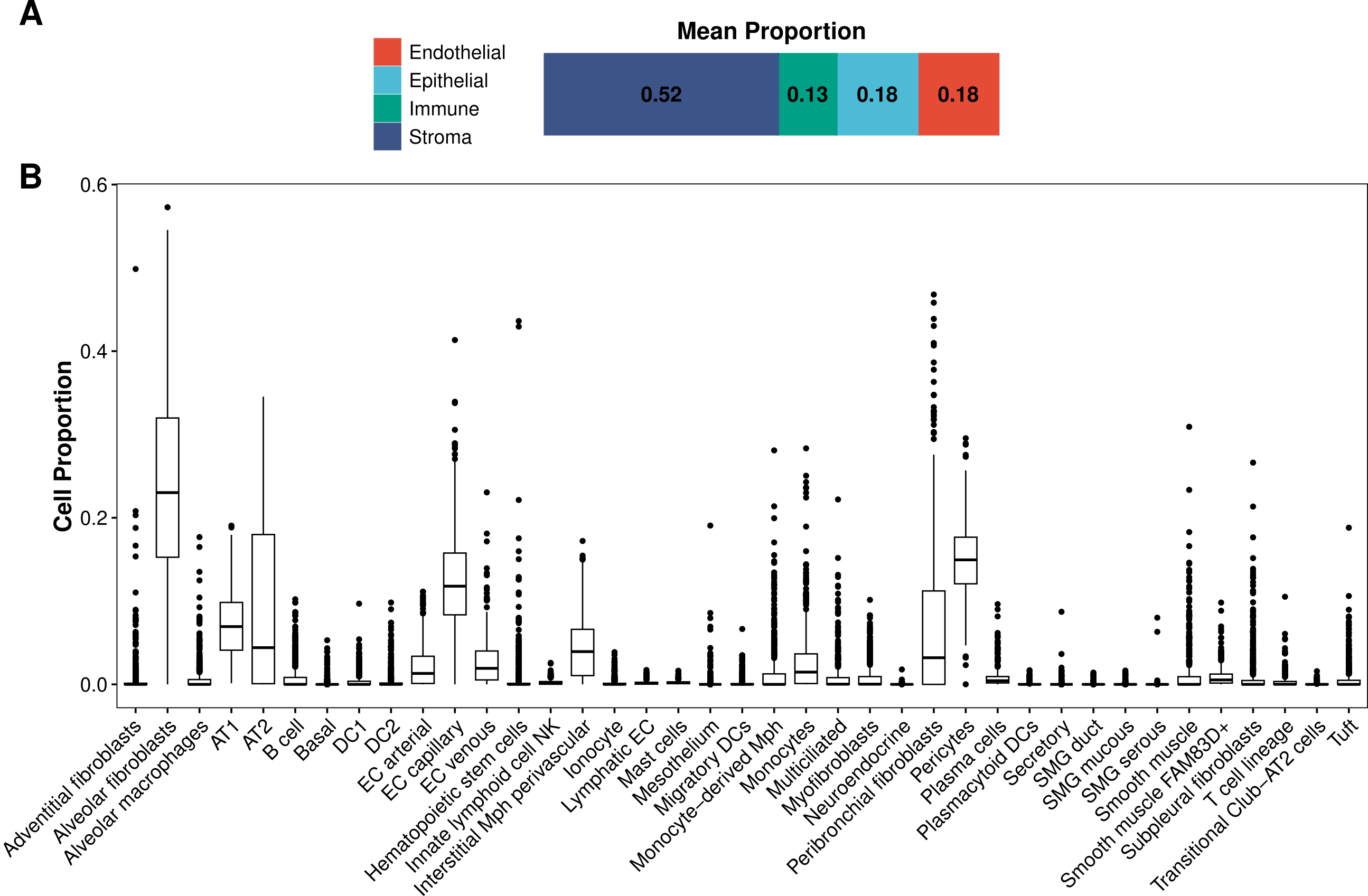

**Figure S5: Cell type deconvolution of the GTEx lung using HLCA v2.** Barplot showing mean cell proportions of **A.** compartment from cell type deconvolution of the GTEx lung. **B.** Box plots showing cell proportion from annotated cell type deconvolution of the GTEx lung (n=578). Box plots show the median and first and third quartiles, whiskers extend to 1.5× the interquartile range, and outlier samples are plotted as black circles.

### **Supplementary Table**

**Table S1: Disease-associated AGTR1 expression across stromal cell populations in the HLCA dataset.** Summary of disease-associated *AGTR1* expression across five *AGTR1*-enriched stromal cell populations in the HLCA dataset. For each cell type, donor-level mean *AGTR1* expression was compared across disease states using Kruskal–Wallis tests, followed by Dunn’s post hoc comparisons with false discovery rate (FDR) correction. Reported values include test statistics, nominal and FDR-adjusted *p* values, and effect size estimates for significant and trend-level comparisons.

**Table S2: Disease-associated AGTR1 expression across pericyte subclusters in the HLCA dataset.** Donor-level comparisons of *AGTR1* expression across transcriptionally defined pericyte subclusters in the HLCA dataset. For each subcluster, disease-associated differences were assessed using Kruskal–Wallis tests, followed by Dunn’s post hoc comparisons where applicable. Reported values include test statistics, nominal and FDR-adjusted *p* values, and effect size estimates for significant and trend-level comparisons.

**Table S3: Disease-associated AGTR1 expression across harmonized pericyte subclusters in a secondary COPD/IPF-enriched dataset.** Analysis of disease-associated *AGTR1* expression across pericyte subclusters HLCA-derived annotations in an independent single-nucleus lung dataset with COPD and IPF donors (control [n=28], COPD [n=18], or IPF [n=32]). Donor-level mean *AGTR1* expression was compared across disease groups using Kruskal–Wallis tests and Dunn’s post hoc comparisons with FDR correction. Reported values include test statistics, nominal and FDR-adjusted *p* values, and effect size estimates for significant and trend-level comparisons.
